# Transplantation of active mitochondria condensed in liquid−liquid phase-separated hydrogels ameliorates myocardial ischemia-reperfusion injury

**DOI:** 10.1101/2025.10.21.683633

**Authors:** Jiacong Ai, Yingxian Xiao, Qishan Li, Yafang Xiao, Weirun Li, Junyao Deng, Weichao Ding, Rui Zhang, Shushan Mo, Yan Zeng, Xueyi Wang, Zhenhua Li

## Abstract

Mitochondrial dysfunction is a key driver of myocardial ischemia-reperfusion injury (MIRI), which exacerbates cardiac damage following the restoration of blood flow. Although single-phase hydrogel-based mitochondria transplantation therapy (MTT) holds promise for restoring cellular energy metabolism, its effectiveness is limited by extracellular calcium-induced mitochondrial degradation. In this study, we design a thermo-sensitive hydrogel utilizing liquid-liquid phase separation (LLPS), composed of gelatin and polyethylene glycol (PEG), to condense freshly isolated mitochondria. Compared to conventional single-phase hydrogels, the LLPS hydrogel significantly increases the sol-gel transition temperature above physiological levels, allowing it to remain injectable at body temperature while enabling rapid mitochondrial release following transplantation. More importantly, the LLPS structure enhances mitochondrial packing density and protects the activity of condensed mitochondria against calcium toxicity through spatial confinement and gelatin-mediated calcium chelation. In vitro, LLPS hydrogel-condensed mitochondria exhibit improved mitochondrial membrane potential and ATP production ability compared to those delivered via single-phase gelatin hydrogel. In vivo, mitochondria released from the LLPS hydrogel are effectively internalized by cardiomyocytes, resulting in improved cardiac function and reduced tissue damage after MIRI. These findings suggest that LLPS-based hydrogels offer a promising strategy to enhance MTT efficacy for MIRI and other mitochondria-related disorders.

## Introduction

Cardiovascular diseases remain the primary cause of mortality worldwide, with ischemic cardiomyopathy representing the most severe and life-threatening manifestation (*1, 2*). While pharmacological thrombolysis and percutaneous coronary intervention (PCI) are necessary means to salvage ischemic myocardium, the sudden reperfusion of blood after ischemia would trigger a cascade of damage processes that may lead to further necrosis (*3–5*). During myocardial ischemia-reperfusion injury (MIRI), pathological Ca^2+^ influx results in calcium overload and elevated reactive oxygen species (ROS) production, subsequently inducing permeability transition pore (mPTP) opening and ultimately precipitating mitochondrial impairment (*6, 7*). Given that mitochondria (Mito) are the primary place of oxidative phosphorylation (OXPHOS) and ATP synthesis, mitochondrial dysfunction leads to a decline in cellular energy metabolism levels (*8–10*). This impairment compromises cellular energetics, disrupts excitation-contraction coupling and promotes maladaptive cardiac remodeling, ultimately contributing to the onset and advancement of heart failure (HF) (*11, 12*).

Current standard pharmacological therapies for MIRI primarily comprise angiotensin-converting enzyme inhibitors (ACEIs), β-blockers, angiotensin II receptor blockers (ARBs) and statins. These agents exert cardioprotective effects predominantly by attenuating adverse ventricular remodeling, largely through reducing cardiac workload, myocardial oxygen demand, and overall myocardial energy requirements (*13, 14*). Despite refinements in coronary reperfusion strategies and optimized clinical management, post-I/R HF mortality and morbidity rates remain unacceptably high (*15, 16*). A fundamental limitation of these therapies lies in their inability to address the underlying energetic deficits of the failing myocardium, rendering them essentially noncurative (*17*). Mitochondria transplantation therapy (MTT) has recently emerged as an innovative approach to restore bioenergetic function in damaged tissues (*18, 19*). In this strategy, viable exogenous mitochondria are introduced into injured tissues, where they are internalized by recipient cells and integrated into the endogenous mitochondrial network (*20*). Transplanted mitochondria have been demonstrated to restore ATP production, regulate mitochondrial dynamics (fusion and fission), enhance mitophagy and autophagy, and modulate the metabolic and inflammatory state of recipient cells (*17, 21–24*). These benefits have been validated in preclinical models of myocardial infarction and pediatric patients with refractory cardiogenic shock, as demonstrated by McCully and colleagues (*22, 25*). In addition, developing strategies to protect mitochondria during transplantation has become a critical research focus (*26, 27*). Recently, some reports have explored several single-phase hydrogel-based delivery systems to enhance mitochondria survival during transplantation(*28*), including thermo-sensitive F127 hydrogels (*29*), alginate-based hydrogels (*30*), and HA-MC hydrogels (*31*).

However, despite the promising potential of MTT, several translational barriers remain. A critical limitation is the vulnerability of isolated mitochondria in the extracellular space, particularly under conditions mimicking the in vivo environment. Extracellular fluids contain approximately 1.8 mmol/L Ca^2+^, a concentration vastly exceeding the physiological intracellular range (50-600 nmol/L) (*32–34*). This high calcium concentration induces mitochondrial swelling, loss of membrane potential, and enzymatic inactivation, significantly reducing the viability and therapeutic efficacy of transplanted mitochondria (*35*). Even when mitochondria are encapsulated in single-phase hydrogels, their homogeneous distribution and the hydrogel’s large pore size allow unrestricted Ca^2+^ exchange, which can still lead to mitochondrial damage. Therefore, preserving mitochondrial activity during delivery and ensuring their functional integration into recipient cells remain major challenges (*36*).

To address this, we develop a mitochondrial condensate system within the sub-compartments of thermo-sensitive hydrogel via liquid-liquid phase separation (LLPS). Simple mixing of polyethylene glycol (PEG), gelatin and freshly isolated mitochondria induces LLPS to form multicompartmentalized hydrogels where mitochondria are condensed in the spherical PEG phase (*37, 38*). Gelatin as the continuous phase endows the LLPS hydrogel with the thermo-sensitive property. Moreover, we find that LLPS significantly alters the sol−gel transition behavior of hydrogels, notably elevating the sol-gel transition temperature to above body temperature (37 °C) (*39*). Remarkably, the condensates of mitochondria located within PEG microdomains increase mitochondrial packing density and improve the activity of mitochondria due to the confinement effect and mitochondria crowding effect. Moreover, the continuous phase of gelatin provides a protective microenvironment which resists the diffusion of Ca^2+^ into the mitochondrial condensates within PEG microdomains via the chelation between the carboxyl group of gelatin and Ca^2+^. This spatial buffering ability to Ca^2+^ mitigates Ca^2+^-induced mitochondrial damage, as confirmed by preserved mitochondria membrane potential (MMP) and ATP production capacity (*40*). Furthermore, both cellular and animal studies demonstrate that the LLPS hydrogel containing condensed mitochondria (Mito@LLPS hydrogel) delivers more functionally active mitochondria to the ischemic myocardium and enhances MTT efficacy compared to single-phase hydrogel-based mitochondria transplantation (Mito@gelatin). More active mitochondria upregulate the mitochondrial fusion protein MFN1 and enhance aerobic respiration while suppressing anaerobic pathways, thereby significantly improving the overall energy metabolism of cardiomyocytes to meet the increased energy demands following MIRI. Thus, by preserving mitochondrial viability, enhancing bioactivity, and enabling controlled delivery to the ischemic myocardium, the Mito@LLPS hydrogel effectively overcomes key limitations of conventional single-phase hydrogel-based MTT strategies. This strategy represents a significant advancement toward the effective clinical translation of mitochondria transplantation for the treatment of MIRI.

## Results

### Preparation and characterization of Mito@LLPS hydrogel

Gelatin, a natural polymer primarily derived from the hydrolysis of collagen, and PEG, a widely utilized biomaterial, both exhibit excellent biocompatibility. Owing to their favorable physicochemical and biological properties, these materials have been extensively employed in translational medicine and biomedical engineering applications. It has been reported that the mixing of PEG and gelatin could induce LLPS (*37*). As a thermo-sensitive polymer, gelatin exhibits a gel state at room temperature (RT) and turns into a sol state at body temperature (37 °C). In contrast, the introduction of PEG not only induced LLPS but also altered the thermal properties of the mixture. The resulting PEG/gelatin formulation remained in a gel state at both RT and 37 °C, indicating an elevated sol-gel transition temperature (Fig. 2A and S1A). Rheological characterization of the PEG/gelatin composite under physiological temperature conditions revealed concentration-dependent viscoelastic behavior. The results demonstrated that both elastic modulus (G′) and viscous (G″) moduli exhibited significant enhancement at 37°C with progressive PEG incorporation (Fig. 2A and 2F) (*41*). Furthermore, the rheological thermomechanical properties of gelatin and PEG/gelatin were affected by their respective concentrations and were systematically investigated using a rheometer across a range of gelatin concentrations (10-25 wt%). Rheological analyses revealed that increasing the gelatin concentration led to a progressive elevation in the sol-gel transition temperature in both single-phase and phase-separated hydrogel systems (Fig. S2A and 2F). Moreover, independent of gelatin concentration, the incorporation of PEG further elevated sol-gel transition temperature (Fig. S2B). Rheological characterization revealed a sol-gel transition temperature of 37°C for homogeneous gelatin solutions at 18.2 wt% concentration. In contrast, the phase-separated hydrogel exhibited a significantly elevated sol-gel transition temperature (>42 °C), which was associated with enhanced overall rheological and mechanical properties (Fig. S2A-B and 2E). The elevated sol-gel transition temperature induced by LLPS endowed the hydrogel with the gel status upon injection into the body, thereby enhancing its retention within the target tissue. Considering both injectability and thermo-mechanical properties, a gelatin concentration (18.2 wt%) was selected for the preparation of the LLPS hydrogel used in all subsequent experiments.

**Fig. 1.**
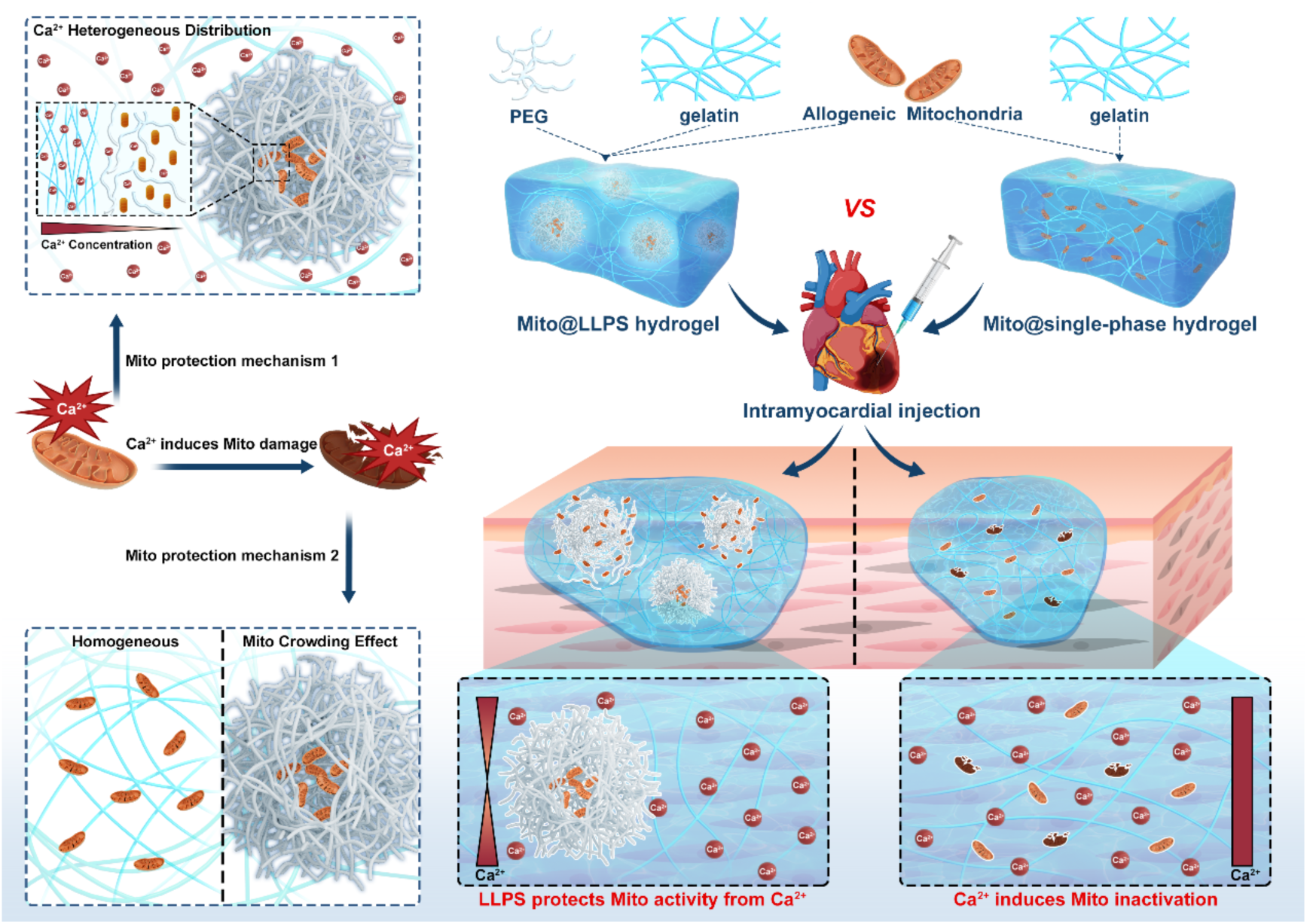
Schematic diagram. Schematic illustrating the mitochondria transplantation therapy (MTT) in myocardial ischemia-reperfusion injury (MIRI) via liquid−liquid phase-separated hydrogel (Mito@LLPS hydrogel) and conventional single-phase hydrogel (Mito@gelatin), respectively.

**Fig. 2.**
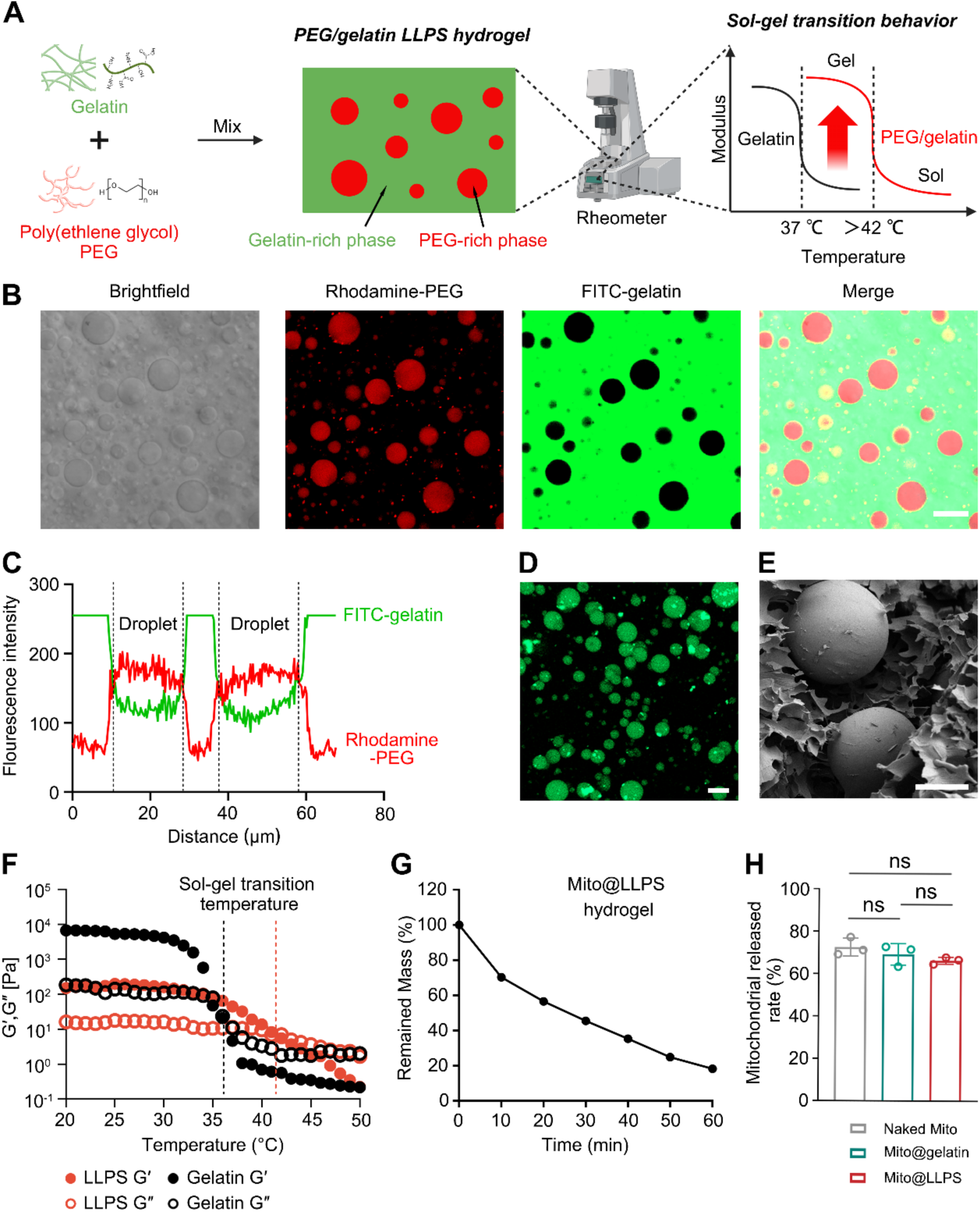
Characterization of Mito@LLPS hydrogel. (A) The schematic illustration of the preparation and characterization of Mito@LLPS hydrogel and sol-gel transition behavior. (B) LCSM image of Mito@LLPS hydrogel with Rhodamine B-labelled PEG and FITC-labelled gelatin. Scale bar: 20 μm. (C) Line-scan profile of the dashed line shown in the panel. (D) LCSM image of Mito_Green_@LLPS hydrogel. The isolated mitochondria were labelled with MitoTracker Green. Scale bar: 20 μm. (E) SEM image of Mito@LLPS hydrogel. PEG appeared as spherical dense structures, and the gelatin phase maintained a continuous, porous network. Scale bar: 20 μm. (F) Rheological properties of Mito@LLPS hydrogel with temperature as variates. PEG (33.3 wt%), gelatin (18.2 wt%) (G) The results of the Mito@LLPS hydrogel disintegration experiment within 1 hour at 37 °C (*n* = 3). (H) The mitochondria release from the Mito@LLPS hydrogel after 3 hours at 37 °C (*n* = 3). Data were presented as mean ± s.d. Statistical significances were calculated via two-tailed unpaired one-way ANOVA-Tukey’s multiple comparisons test.

Then, freshly isolated mitochondria were introduced into this LLPS system, and a mitochondria-condensed LLPS hydrogel (Mito@LLPS hydrogel) was prepared by mixing PEG solution, gelatin solution and freshly isolated mitochondria, followed by vortexing at 37 °C (Fig. 2A). Laser scanning confocal microscope (LSCM) images of Rhodamine B-labelled PEG and FITC-labelled gelatin showed that gelatin (18.2 wt%) and PEG (33.3 wt%) constituted LLPS structures to form PEG/gelatin LLPS hydrogel at 25 °C (Fig. 2B). The line scan images demonstrated that the PEG/gelatin LLPS hydrogel consisted of gelatin-rich dispersed phase and PEG-rich droplet phase (Fig. 2C). Mitochondria were isolated from the hearts of healthy SD rats by differential centrifugation. Then, the purity (99.85%) of the extracted mitochondria was confirmed by a flow cytometer with MitoTracker Green-labelled isolated mitochondria (Mito_Green_) (Fig. S3A). To determine the spatial distribution of mitochondria within the hydrogel, Mito_Green_ were also employed. The image of LSCM revealed that mitochondria were condensed within the PEG-rich phase (Fig. 2D). Further structural characterization using scanning electron microscopy (SEM) demonstrated that the PEG phase exhibited a dense, non-porous morphology, whereas the gelatin phase was loose and porous (Fig. S4). SEM images of the Mito@LLPS hydrogel (Fig. 2E) revealed a distinct biphasic architecture. PEG formed discrete, densely packed spherical domains, indicative of the condensed LLPS phase. In contrast, the gelatin component constituted a continuous and porous network, as further corroborated by supplementary images (Fig. S4). This microstructural organization supported the phase-separated nature of the LLPS hydrogel and highlighted the spatial segregation of its components. Notably, mitochondria were not directly visible in the SEM images due to their encapsulation within the non-porous PEG matrix. To mimic the condition of LLPS hydrogels with tissue fluid in vivo, LLPS hydrogels were immersed in PBS, and hydrogel desolvation experiments were performed. Notably, the Mito@LLPS hydrogel exhibited significant structural instability, undergoing nearly complete degradation within 1 hour at 37 °C (Fig. 2G). Moreover, to investigate the mitochondrial release behavior from the Mito@LLPS hydrogel, release studies were conducted using a Transwell system (*29*). Quantitative analysis of mitochondrial protein content in the lower chamber of a Transwell system revealed that 66.32% of mitochondria condensed within the Mito@LLPS hydrogel were released into the lower compartment within 1 hour at 37 °C. Notably, this release efficiency was statistically comparable to that of bare mitochondria, indicating that the LLPS hydrogel did not hinder mitochondrial release, mimicking the in vivo conditions (Fig. 2H). Moreover, employing the same analytical approach, the release profile of mitochondria from the Mito@LLPS hydrogel was evaluated following ultraviolet (UV) light-induced crosslinking of gelatin methacryloyl (GelMA). It was found that after chemical crosslinking for 5 or 10 seconds, the chemical crosslinked Mito@LLPS hydrogel only released 12.23% and 12.17% of mitochondria at 37 °C in 1 hour, respectively. These results demonstrated that chemical crosslinking within the Mito@LLPS hydrogel significantly restricted mitochondrial release, thereby hindering its suitability for mitochondria transplantation (Fig. S5). In contrast, the non-crosslinked LLPS hydrogel presented a more permissive network structure, offering minimal resistance to mitochondrial release. Collectively, these findings indicated that the Mito@LLPS hydrogel not only effectively condensed mitochondria but also enabled their rapid release in a PBS environment, making it a promising vehicle for mitochondria transplantation.

### Mitochondria protection by LLPS hydrogel

As designed, the Mito@LLPS hydrogel condensed mitochondria in the PEG phase as the subcompartment of the hydrogel, which provided a compact and restricted microenvironment for mitochondria (Fig. 3A). To evaluate the protective effect of Mito@LLPS hydrogel on mitochondrial activity and functions, mitochondria were co-cultured with 1.8 mmol/L Ca^2+^ for 1 hour under three conditions: as naked mitochondria, dispersed in gelatin hydrogel, and condensed in LLPS hydrogel. Following incubation, MMP and ATP production were assessed, respectively. Among the groups, the Mito@LLPS hydrogel exhibited the most effective protection, as evidenced by higher MMP and ATP levels compared to both the naked Mito and Mito@gelatin groups (Fig. 3B-C).

**Fig. 3.**
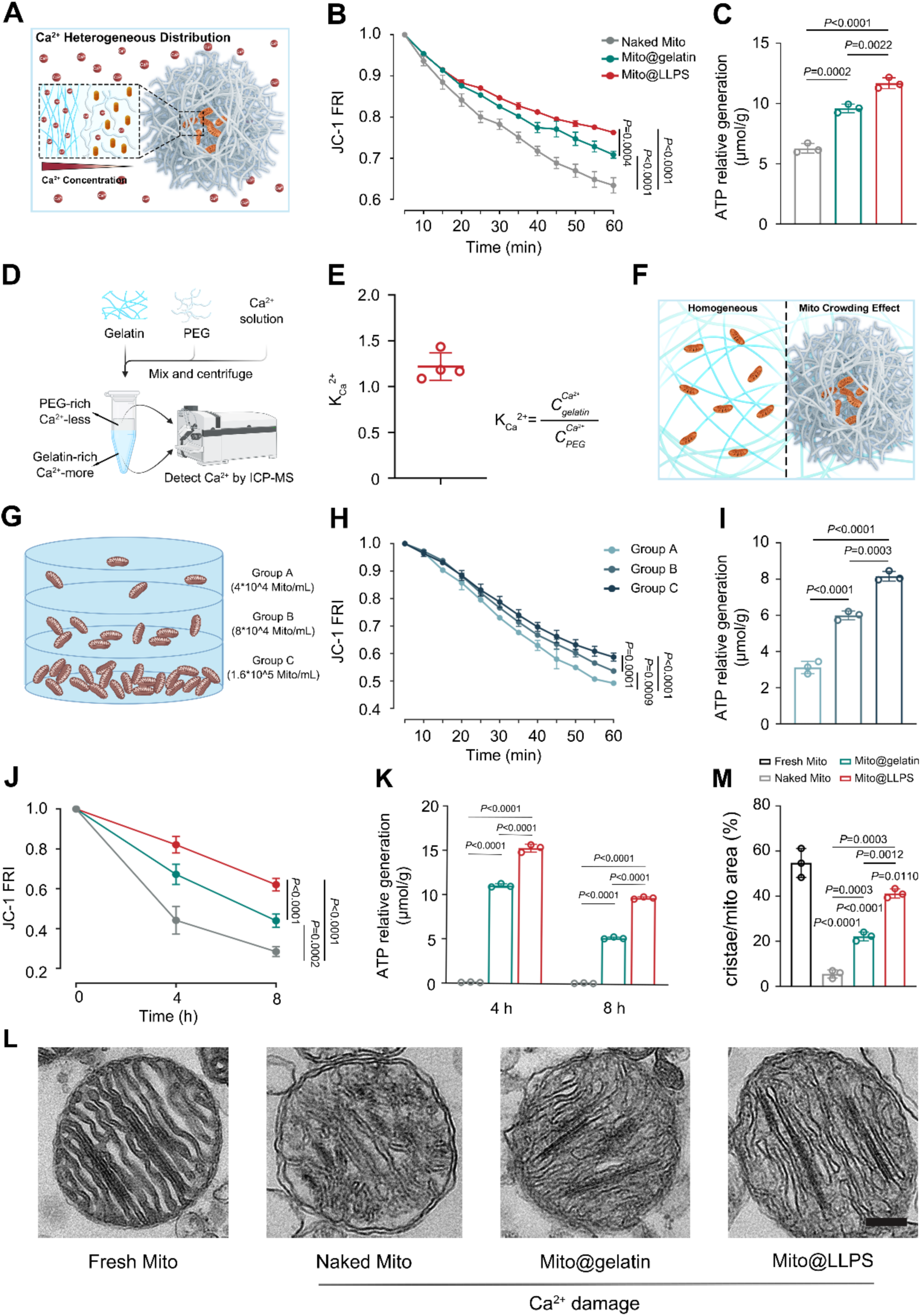
Mitochondria protection by liquid−liquid phase separated hydrogel. (A) Schematic illustration of the Ca^2+^ heterogeneous distribution in Mito@LLPS hydrogel. (B) Time-series quantitative analysis of JC-1 staining as an indicator of MMP after Ca^2+^ treatment (*n*=3). (C) Results of mitochondrial ATP relative generation after Ca^2+^ treatment (*n*=3). (D) Schematic illustration of the process of Ca^2+^ concentration detection in two separated phases by ICP-MS. (E) The calcium distribution coefficient which was calculated by the ratio of the concentration of Ca^2+^ in the gelatin phase to the concentration in the PEG phase measured by ICP-MS (*n*=4). (F) Schematic illustration of Mito crowding phenomenon in Mito@LLPS hydrogel contrast with Mito@gelatin hydrogel. (G) Schematic illustration of an experimental model with different Mito crowding (group A-C) to investigate the Mito crowding effects on Mito activity. (H) Time-series quantitative analysis of JC-1 staining as an indicator of MMP in group A-C (*n*=3). (I) Results of mitochondrial ATP relative generation per mitochondria in group A-C (*n*=3). (J) Time-series quantitative analysis of JC-1 staining as an indicator of MMP at 4 °C. (K) Results of mitochondrial ATP relative generation at 4 °C (*n*=3). (L) TEM image of isolated mitochondria ultrathin section (scale bar, 100 nm). (M) Statistical data for the ratio of mitochondria inner membrane cristae area to mitochondria cross-sectional area (*n*=3). Data were presented as mean ± s.d. For (B), (H), (J), (K), Statistical significances were performed via two-tail two-way ANOVA-Bonferroni’s multiple comparisons test. For (C), (I), (M), Statistical significances were performed via two-tail one-way ANOVA-Bonferroni’s multiple comparisons test.

As one underlying mechanism for the mitochondria protection effect, the presence of chelation between the carboxyl group of gelatin and Ca^2+^ resulted in a concentration gradient of Ca^2+^ between the PEG and gelatin phases within the Mito@LLPS hydrogel (Fig. 3A). To better exhibit the different distribution of Ca^2+^ in the two phases, the distribution coefficient of Ca^2+^ was investigated, defined as the concentration ratio of Ca^2+^ between the gelatin and PEG phases at equilibrium. The calculation formula of the partition coefficient of Ca^2+^ in PEG/gelatin phase-separated solution was as follows: 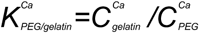. The separation between gelatin and PEG phases was accelerated by centrifugation at a Ca^2+^ concentration of 1.8 mmol/L. Subsequently, inductively coupled plasma-mass spectrometry (ICP-MS) analysis quantified the differential Ca^2+^ distribution across phases at equal volumetric sampling (Fig. 3D). As the result, the distribution coefficient 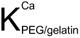 was calculated as 1.22, indicating a higher concentration of Ca^2+^ in the gelatin phase compared to the PEG phase. This finding suggested a heterogeneous distribution of Ca^2+^ within the PEG/gelatin LLPS hydrogel, with resistance to Ca^2+^ accumulation within the mitochondria-condensed PEG microdomains (Fig. 3E).

Furthermore, as the other underlying mechanism for the mitochondria protection effect, the spatial confinement of mitochondria in PEG microdomains reduced the distance between mitochondria and enhanced mitochondria crowding. Such confinement effect of the LLPS hydrogel and the mitochondria crowding effect promoted the accumulation of mitochondrial membranes, thereby reducing the density of calcium ions per unit area of the mitochondrial membrane (Fig. 3F). As a result, Ca^2+^-induced mitochondrial damage was alleviated, and mitochondrial activity was enhanced. To confirm it, MMP and ATP production capacity were measured to investigate the relation between mitochondrial activity and the mitochondria crowding effect. Mitochondria were dispersed at three concentration gradients (group A: 4×10^4^ Mito/mL; group B: 8×10^4^ Mito/mL; group C:1.6×10^5^ Mito/mL) in PEG solution (Fig. 3G). The results exhibited that the higher concentration of mitochondria induced the higher resistance to Ca^2+^ invasion, along with enhanced MMP and ATP production capacity per mitochondria (Fig. 3H-I).

Therefore, by combining these two factors, the Mito@LLPS hydrogel exhibited the ability of Ca^2+^ buffering, which effectively resisted Ca^2+^-induced mitochondrial damage and protected mitochondrial activity. Moreover, the potential of the Mito@LLPS hydrogel to preserve mitochondrial functions during low-temperature storage was also evaluated. Naked Mito, Mito@gelatin and Mito@LLPS hydrogel were stored at 4 °C. At predetermined time points (0, 4, and 8 hours), mitochondria were collected from each group, and their MMP and ATP levels were subsequently quantified to assess mitochondrial activity over time. Compared to both naked mitochondria and the Mito@gelatin group, the Mito@LLPS hydrogel demonstrated a superior capacity to preserve mitochondrial function under low-temperature conditions, as evidenced by sustained MMP and ATP levels (Fig. 3J–K).

Furthermore, mitochondrial morphology across different experimental groups was also examined by transmission electron microscopy (TEM). Freshly isolated mitochondria and the mitochondria from each treatment groups of naked Mito, Mito@gelatin, Mito@LLPS hydrogel after Ca^2+^ treatment were collected. Then the ultrathin sections were used to observe the morphological changes of the mitochondria inner membrane cristae by TEM (*42*). Notably, mitochondria from the Mito@LLPS hydrogel retained a more normal cristae morphology compared to other groups (Fig. 3L-M). These results suggested that compared with the traditional single-phase hydrogel, the LLPS hydrogel offered enhanced protection of mitochondria against Ca^2+^-induced damage and supported the maintenance of mitochondrial activity and functions under storage conditions.

### The mitochondria transfer from Mito@LLPS hydrogel to recipient cells

In vivo imaging system (IVIS) experiments were performed using MitoTracker Deep Red FM-labelled mitochondria (Mito_Red_) to evaluate their retention and distribution following transplantation. Both Mito_Red_@LLPS hydrogel and free Mito_Red_ were administered through intramyocardial injection in rats’ ventricles. Subsequent IVIS imaging of the major organs (heart, liver, spleen, lung, kidneys) was performed at predetermined intervals (6, 12, 24, and 48 hours) post-administration. The quantitative fluorescence analysis revealed significantly greater mitochondrial retention in the Mito@LLPS group compared to naked Mito at all timepoints (p<0.05), with 2.3-fold higher mean intensity at 48 hours post-transplantation (Fig. 4A-B). The results revealed that mitochondria condensed in LLPS hydrogel exhibited significantly prolonged retention in cardiac tissue compared to naked mitochondria. Additionally, it can be observed that the fluorescence intensity of the liver and kidneys was enhanced after intramyocardial injection of Mito_Red_. This indicated that mitochondria those not internalized by cardiac cells were predominantly metabolized and cleared through the liver and kidneys, similar to the metabolic pathway of most macromolecular drugs (Fig. 4C). Moreover, Cy5-labelled gelatin was also used to evaluate the retention of the Mito@LLPS hydrogel. To evaluate its biodistribution and retention, major organs were collected at 0, 12, 24, and 48 hours post-injection and analyzed using ex vivo fluorescence imaging. The IVIS images showed that the Mito@LLPS hydrogel was quickly metabolized after being injected into the heart, and the main metabolic sites remained the liver and kidneys (Fig. S6).

**Fig. 4.**
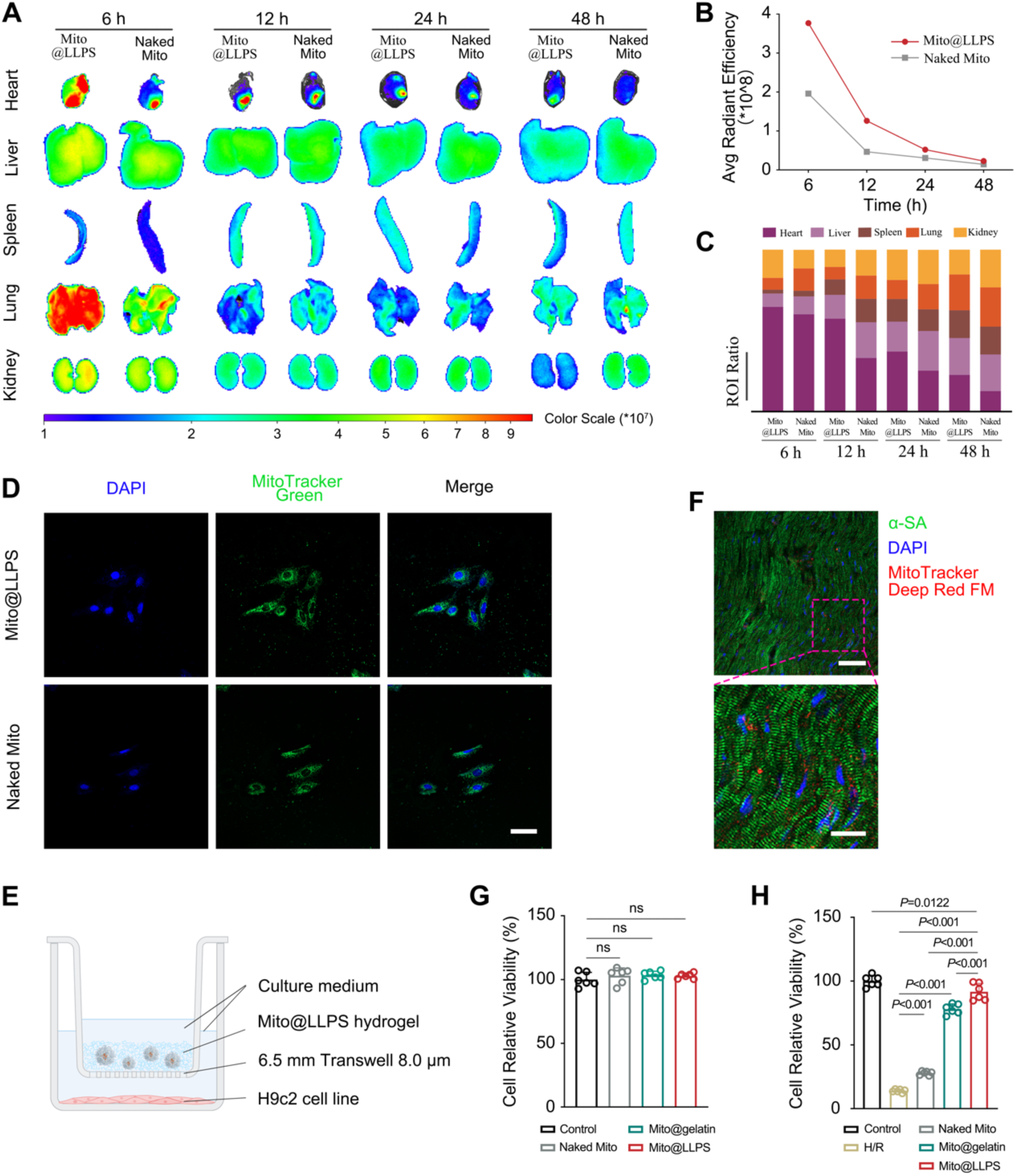
The mitochondria transfer from Mito@LLPS hydrogel to recipient cells. (A) IVIS imaging of major organs (heart, liver, spleen, lung, kidney) collected from rats at 6 h, 12 h, 24 h and 48 h post-intramyocardial injection of naked Mito and Mito@LLPS hydrogel. All mitochondria were labelled with MitoTracker Deep Red FM. (B) Quantitative analysis of average radiant efficiency of hearts at 6-h, 12-h, 24-h and 48-h post-intramyocardial injection of naked Mito and Mito@LLPS hydrogel. (C) Relative quantitative analysis of average radiant efficiency of major organs at 6 h, 12 h, 24 h and 48 h post-intramyocardial injection of naked Mito and Mito@LLPS hydrogel. (D) LCSM image of cellular uptake of isolated mitochondria in naked Mito, Mito@gelatin and Mito@LLPS hydrogel groups. Cell nuclei were stained with DAPI (blue), and isolated mitochondria were stained using MitoTracker Green (green). Scale bars, 20 μm. (E) Schematic illustration of cell experiments conducted using a Transwell system. (F) LCSM images of cardiac cells after the uptake of Mito@LLPS hydrogel. Cell nuclei were stained with DAPI (blue), and isolated mitochondria were visualized using MitoTracker Deep Red FM (red). Scale bars, top: 50 μm, bottom: 20 μm. (G-H) CCK-8 assay results demonstrating the relative viability of cardiomyocytes following incubation with PBS (control), naked Mito, Mito@gelatin, Mito@LLPS hydrogel, under normoxic (G) and hypoxia-reoxygenation (H/R) (H) conditions (*n*=6). Statistical significance was calculated via two-tail one-way ANOVA-Tukey’s multiple comparisons test.

To investigate the release and cellular uptake of mitochondria from Mito@LLPS hydrogel, in vitro Transwell assays were performed. Mitochondria were labelled with MitoTracker Green and incorporated into various formulations, including naked Mito_Green_, Mito_Green_@gelatin, and Mito_Green_@LLPS hydrogel. These formulations were applied to the upper chamber of Transwell permeable supports, while H9c2 cardiomyocytes were seeded in the lower chamber (Fig. 4E). CLSM confirmed the successful uptake of mitochondria, released from the upper compartments, by cardiomyocytes. Comparable fluorescence intensities corresponding to the transplanted mitochondria were observed within cardiomyocytes across the naked Mito, Mito@gelatin, and Mito@LLPS groups, indicating equivalent levels of mitochondrial internalization among these groups (Fig. 4D, S7). Furthermore, to confirm the uptake of mitochondria by cardiac cells in vivo, mitochondria were labelled with MitoTracker Deep Red FM, and Mito_Red_@LLPS hydrogel was injected into the ventricular muscle of rats via intramyocardial injection. Then, the rats were sacrificed three hours post-injection, and the hearts were collected, sectioned, and subjected to immunofluorescent staining with α-SA (green) and DAPI (blue). LSCM analysis indicated efficient internalization of released mitochondria from Mito_Red_@LLPS hydrogel by cardiomyocytes within a short time frame (Fig. 4F). These findings collectively demonstrated that mitochondria condensed in LLPS hydrogel were capable of rapid release and efficient uptake by cardiac cells in both cell and animal models.

Moreover, CCK-8 were performed to investigate the therapeutic effect of mitochondria transplantation on cardiomyocytes. No significant difference was observed in cell proliferation among the naked Mito, Mito@gelatin, and Mito@LLPS hydrogel groups compared to the control, suggesting excellent biocompatibility and biosafety of the Mito@LLPS hydrogel formulation (Fig. 4G). In contrast, under hypoxia-reoxygenation (H/R) conditions, the Mito@LLPS hydrogel group demonstrated superior therapeutic effects relative to both naked Mito and Mito@gelatin groups, with the naked Mito group showing only modest efficacy (Fig. 4H). Furthermore, the in vivo biosafety experiment of Mito@LLPS hydrogel was performed following intramyocardial injection. Echocardiography measurements indicated no adverse effects on left ventricular ejection fraction (LVEF) or fractional shortening (LVFS), and cardiac function remained unaltered seven days post-injection (Fig. S8). Histological evaluation was also performed on major organ systems (heart, liver, spleen, lung, and kidney) using hematoxylin and eosin (H&E) staining. Tissue sections from each experimental group were examined for structural integrity and pathological alterations (Fig. S9). Thus, these results confirmed that Mito@LLPS hydrogel was a histocompatible and biocompatible delivery system, suitable for MTT, without eliciting adverse effects in healthy cardiac tissue. Moreover, these results also confirmed that mitochondrial transplantation has no proliferation and function promotion on normal cardiomyocytes or individuals.

### Therapeutic effects of Mito@LLPS hydrogel in vitro

All cellular experiments were performed using a Transwell system. Naked Mito, Mito@gelatin and Mito@LLPS hydrogel were placed in the upper compartment, while cardiomyocytes were seeded in the lower compartment (Fig. 4E) (*29*). In order to elucidate the impacts induced by MTT on the aerobic or anaerobic metabolic components of cellular energy metabolism, oxygen consumption rate (OCR), ATP real-time production rate and extracellular acidification rate (ECAR) were analyzed. As the results, compared to naked Mito and Mito@gelatin groups, the Mito@LLPS hydrogel significantly enhanced the OCR and ATP production rate of H9c2 cell lines, but reduced ECAR (Fig. 5A, S10A). Quantitative analysis of OCR measurements demonstrated that the Mito@LLPS hydrogel group exhibited superior therapeutic effects on mitochondrial respiration compared to both naked Mito and Mito@gelatin treatment groups. This was evidenced by a significant increase in basal respiration, ATP-linked respiration, maximal respiratory capacity, and spare respiratory capacity (Fig. 5B). Similarly, quantitative analysis of ECAR measurements revealed that MTT via Mito@LLPS hydrogel could reduce the basal glycolytic, glucose metabolism, glycolytic capacity and glycolytic reverse of cells after H/R injury (Fig. S10B). Additionally, the analysis of the real-time ATP production rate confirmed that after MTT, the overall cellular ATP production rate of Mito@LLPS hydrogel group was significantly enhanced compared to other groups (Fig. 5C). Notably, the amount of ATP produced through mitochondrial aerobic respiration significantly increased, while the ATP production rate by anaerobic glycolysis was greatly reduced. Moreover, the ratio of ATP generated via mitochondrial aerobic respiration to that produced through anaerobic glycolysis was markedly decreased (Fig. 5D). Combined with the data of OCR and ECAR, these indicated that Mito@LLPS could promote mitochondrial metabolism in cells, improve anaerobic glycolysis, and significantly enhance the ATP production level of the cells. In conclusion, all the results indicated that MTT via Mito@LLPS hydrogel could promote ATP production from aerobic respiration and decrease anaerobic respiration.

**Fig. 5.**
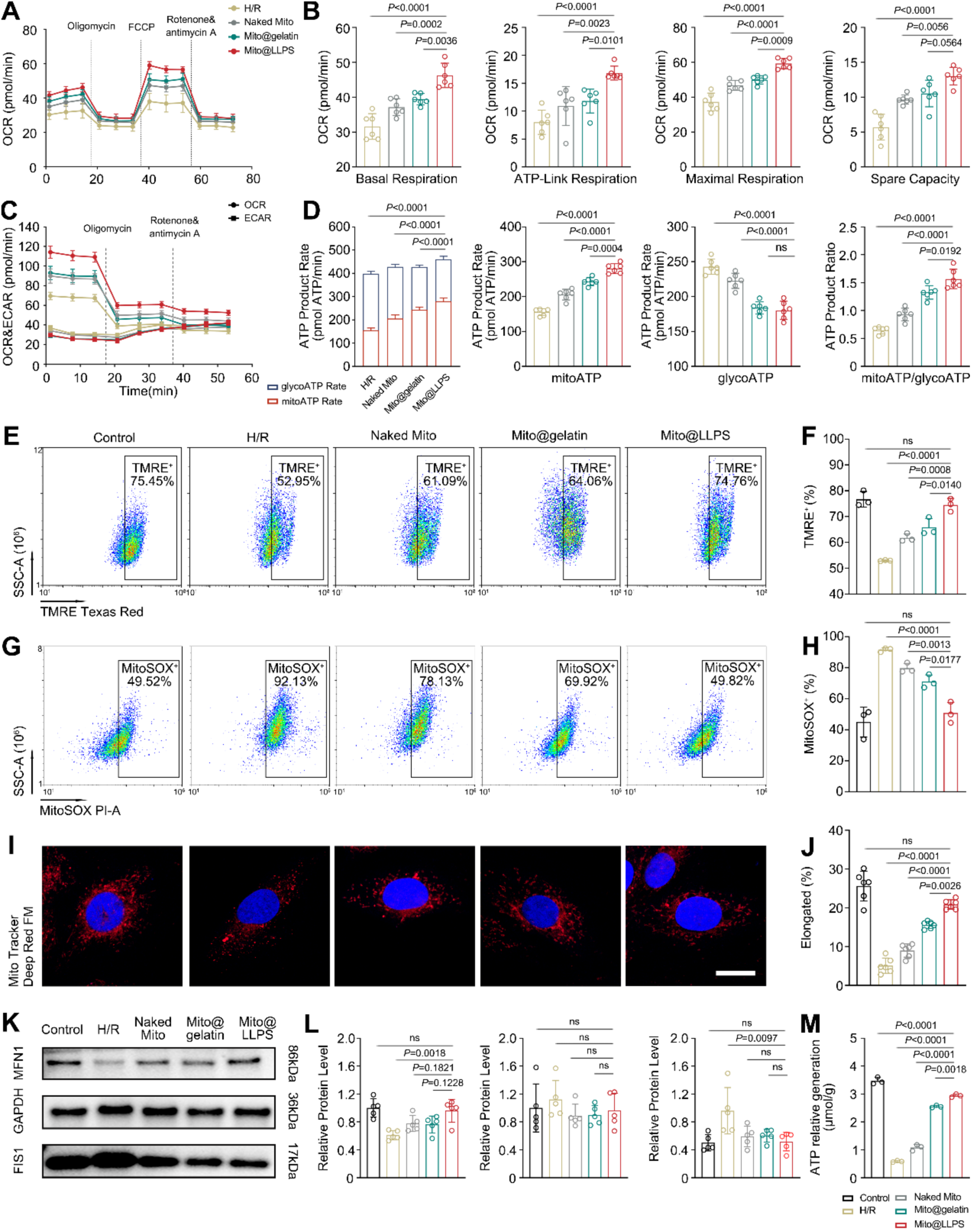
Therapeutic effects of Mito@LLPS hydrogel in vitro. (A-B) The results of cellular OCR (A) and quantitative analyses of OCR (basal respiration, ATP-link respiration, maximal respiration and spare capacity) (B) after MTT treatment of HR-injured cells (*n*=6). (C-D) The results of cellular real-time ATP rate assay (C) and quantitative analyses of cellular real-time ATP rate assay (Total ATP product rate, mitoATP, glycolATP, mitoATP/glycolATP) (D) after different MTT treatments in H/R-injured cardiomyocytes (*n*=6). (E-F) Image of flow cytometry (E) and quantitative analyses (F) of TMRE staining after different MTT treatments in H/R-injured cardiomyocytes (*n*=3). (G-H) Image of flow cytometry (G) and quantitative analyses (H) of TMRE staining after different MTT treatments in H/R-injured cardiomyocytes (*n*=3). (I-J) LCSM images (I) and the responding quantitative analysis (J) of mitochondria network morphology after different MTT treatments in H/R-injured cardiomyocytes (*n*=6). Cell nuclei were stained with DAPI (blue), and mitochondria were visualized using MitoTracker Green (green). Scale bars, 10 μm. (K-L) WB analysis (K) and quantitative analyses (L) of MFN1 and FIS1 expression after different MTT treatments in H/R-injured cardiomyocytes (*n*=5). (M) Total cellular ATP concentration level after different MTT treatments in H/R-injured cardiomyocytes (*n*=3). For (F), (H), (J), (L) and (M), statistical significances were performed via two-tail one-way ANOVA-Tukey’s multiple comparisons test. For (B) and (D), statistical significances were performed via two-tail one-way ANOVA-Bonferroni’s multiple comparisons test.

Furthermore, the therapeutic effect of Mito@LLPS hydrogel was demonstrated by Calcein AM/PI staining. The images of LSCM and quantitative analysis showed that Mito@LLPS hydrogel could effectively improve the survival status of cardiomyocytes (Fig. S11). Flow cytometry with TMRE staining revealed that mitochondria transplantation with Mito@LLPS hydrogel effectively ameliorated the MMP in cardiomyocytes following H/R injury (Fig. 5E-F). Furthermore, MitoSOX red staining results indicated that the MTT with Mito@LLPS hydrogel significantly improved the oxidative stress condition after H/R (Fig. 5G-H). These results indicated that mitochondria transplantation via Mito@LLPS hydrogel could promote the cell activity after H/R treatment, restore MMP and inhibit the production of mitochondrial superoxide (MitoSOX).

To further investigate the cardioprotective effects mechanism of mitochondria transplantation with Mito@LLPS hydrogel, MitoTracker Deep Red FM was used to label transplanted mitochondria. Post-treatment, H/R-injured cardiomyocytes exhibited ovoid, branched mitochondria with a relatively complete network structure following Mito@LLPS hydrogel transplantation (Fig. 5I-J, S12). Finally, the expression levels of mitochondria dynamics-related proteins were assessed by western blot (WB) experiments. The mitochondria transplantation with Mito@LLPS hydrogel increased the expression of fusion-related protein MFN1, while the expression levels of division-related protein FIS1 decreased. This finding indicated that the exogenously supplemented mitochondria could integrate into the mitochondria pool and maintain the mitochondrial homeostasis of the recipient cardiomyocytes (Fig. 5K-L). To assess the whole energy metabolic supply to H/R injured cells after mitochondria transplantation, cellular overall ATP levels were also measured. Notably, cells after H/R treated with Mito@LLPS hydrogel exhibited the highest ATP levels compared to those receiving naked Mito or Mito@gelatin (Fig. 5M).

### Therapeutic effects of Mito@LLPS hydrogel in vivo

To evaluate the cardioprotective effect of Mito@LLPS hydrogel-based MTT in animal models, a rat model of MIRI was constructed. Following MIRI, rats were administered one of the following treatments: PBS, naked Mito, Mito@gelatin, or Mito@LLPS hydrogel. Specifically, these rats underwent a 45-minute left anterior descending artery ligation surgery, and then the ligation band was removed to initiate the reperfusion process (*43*). Then, PBS, LLPS hydrogel, naked Mito, Mito@gelatin, and Mito@LLPS hydrogel were intramyocardially injected into the left ventricular of MIRI rats, respectively (*44*). Twenty-eight days after surgery, the echocardiographic evaluation of cardiac function showed that the LVEF in the I/R group significantly decreased, from 88.05% to 47.95%. In contrast, treatment with Mito@LLPS hydrogel resulted in a markedly improved LVEF of 80.21%, outperforming among LLPS hydrogel (57.21%) naked Mito (59.26%) and Mito@gelatin (66.45%) groups (Fig. 6A-B). Similarly, other functional parameters, including LVFS, left ventricular end-systolic diameter (LVD,S) and left ventricular end-systolic volume (LVV,S), in the Mito@LLPS hydrogel group were more closely aligned with those of the control group, in contrast to other treatment groups (Fig. 6C, S13). Notably, there were no statistically significant differences detected among groups for left ventricular end-diastolic diameter (LVD,D) and left ventricular end-diastolic volume (LVV,D) (Fig. S13).

**Fig. 6.**
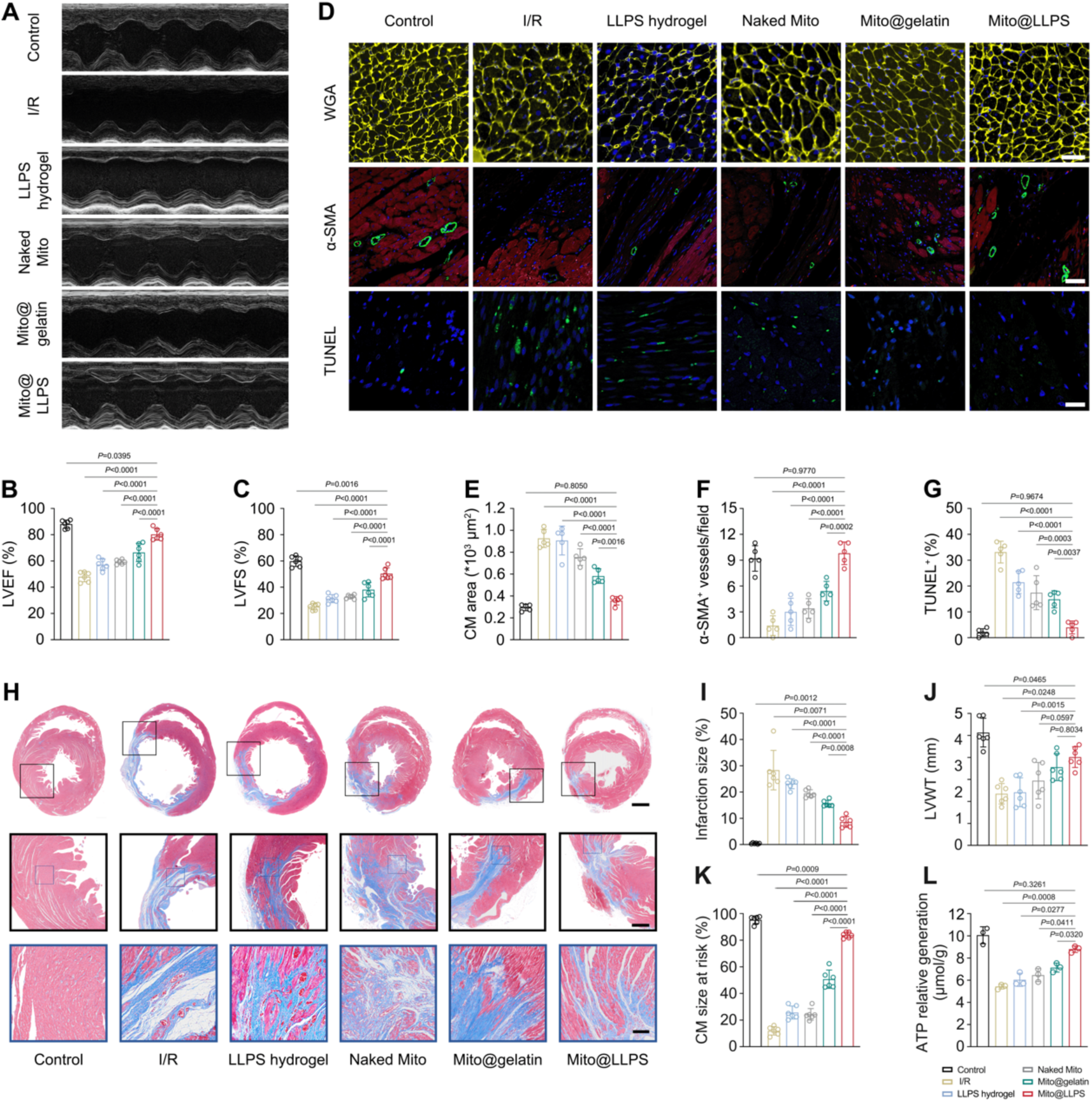
Therapeutic effects of Mito@LLPS hydrogel in vivo. (A-C) Echocardiographic assessment and quantitative analysis of LVEF and LVFS after different MTT treatments on I/R-injured rats (*n*=6). (D-G) Fluorescence images (D) of WGA (E), α-SMA (F) and TUNEL (G) assay and quantitative analysis of cardiac sections from I/R-injured rats after different MTT treatments (*n*=5). Scale bar (WGA), 40 μm; Scale bar (α-SMA), 100 μm; Scale bar (TUNEL), 50 μm. (H) Masson’s trichrome staining images of infarction. Scale bar, top: 2000 μm, middle: 750 μm, bottom: 150 μm. (I-K) The quantitative analysis according to Masson’s trichrome staining images, including the infarction size (I), the thickness of the ventricular infarcted wall (J) and viable myocardium (K) in cardiac sections from I/R-injured rats following different MTT treatments (*n*=6). (L) Quantification of total cellular ATP levels in ventricular tissues from I/R-injured rats following different MTT treatments (*n*=3). Statistical significances were performed via two-tail one-way ANOVA-Tukey’s multiple comparisons test (B-G, J-K) and two-tail one-way ANOVA Brown-Forsythe and Welch multiple comparisons test (I, L).

Next, histological assessments were conducted at designated time points to evaluate therapeutic efficacy. Cardiomyocyte hypertrophy and neovascularization were examined 28 days post-treatment using WGA and α-SMA staining, respectively. Apoptotic cell death within the infarct border zone was quantified at 7 days post-treatment by TUNEL assay (Fig. 6D). These results demonstrated that MTT via Mito@LLPS hydrogel significantly attenuated MIRI-induced compensatory cardiomyocyte hypertrophy, mitigated adverse cardiac remodeling, enhanced angiogenesis, and suppressed cellular apoptosis in the MIRI zone compared to naked Mito and Mito@gelatin groups (Fig. 6E-F). These findings further indicated that Mito@LLPS hydrogel treatment effectively and rapidly repaired MIRI-mediated myocardial damage. By day twenty-eight, rats in the untreated I/R group exhibited significantly larger infarct sizes and more extensive fibrotic scar formation. In contrast, MTT via the Mito@LLPS hydrogel resulted in markedly reduced scar area, increased viable myocardium, and preserved infarct wall thickness compared to LLPS hydrogel, naked Mito and Mito@gelatin groups (Fig. 6H-K). These findings indicated that the healthy mitochondria transplantation in the Mito@LLPS hydrogel group could prevent adverse myocardial remodeling and contribute significantly to the amelioration of cardiac function. In comparison, the plain PEG/gelatin LLPS hydrogel exhibited no significant therapeutic effect on MIRI. Similarly, the H&E staining of myocardial tissue sections revealed that MIRI resulted in cardiac scar formation and atrophic remodeling due to extensive myocardial fibrosis, accompanied by disorganized cardiomyocyte alignment. Following treatment with the Mito@LLPS hydrogel, myocardial fibrosis was markedly reduced, accompanied by a more organized and orderly alignment of cardiomyocytes (Fig. S14). Consistent with these findings, ATP levels in myocardial tissue measured on day 7 post-treatment were significantly elevated in the Mito@LLPS hydrogel group (Fig. 6L). This enhancement in ATP content further supported the superior capacity of the Mito@LLPS system to restore energy metabolism in hearts affected by MIRI.

## Discussion

MIRI is characterized by an abrupt restoration of blood flow that paradoxically exacerbates cardiac tissue damage, largely driven by mitochondrial dysfunction, calcium overload, and oxidative stress. MTT has emerged as a promising therapeutic strategy to replenish damaged cells with functional mitochondria (*45, 46*). However, the adverse transplanted microenvironment of the heart after MIRI—with high concentrations of ROS, Ca^2+^ overload, and pro-inflammatory cytokines—poses a serious threat to the viability and functionality of the transplanted mitochondria (*8, 47, 48*). Despite growing interest in MTT, many early studies overlooked the detrimental impact of the post-I/R microenvironment on transplanted mitochondria, particularly the cytotoxic effects of extracellular and blood Ca^2+^ (*36*). Minimizing mitochondrial damage during transfer is crucial, particularly when donor mitochondria are limited and of high clinical value. This oversight leads to an ongoing debate regarding the true efficacy of mitochondria transplantation. Thus, we aim to investigate whether mitochondria with preserved functionality exert a more pronounced reparative effect on damaged recipient cells compared to dysfunctional mitochondria. We hypothesize that preserving the functional integrity of mitochondria during transplantation is extremely important for the successful functional recovery of injured recipients. To confirm this, a novel LLPS hydrogel with condensed mitochondria (Mito@LLPS hydrogel) is designed, which not only protects mitochondria from Ca^2+^-mediated damage but also enhances their retention and bioenergetic functions. By addressing these fundamental questions, our work aims to advance mitochondria transfer from a promising concept to a clinically viable treatment for MIRI.

The Mito@LLPS hydrogel is formed through LLPS between gelatin and PEG, with mitochondria preferentially partitioning into the PEG-rich phase. This architecture allows the mitochondria to be densely packed within spherical PEG domains, while the surrounding gelatin-rich continuous phase serves as a semi-permeable matrix that buffers against Ca^2+^ influx. Chelation of Ca^2+^ by the carboxyl groups in gelatin, along with spatial segregation of PEG-condensed mitochondria, collectively creates a localized environment with reduced ionic toxicity. In addition, LLPS-mediated condensation of mitochondria not only provides a protective environment but also induces a mitochondrial crowding effect that promotes oxidative phosphorylation (OXPHOS) efficiency (*49, 50*). These results are consistent with the previous research results that confinement and biomacromolecule crowding within biomimetic compartments can enhance enzymatic kinetics and organelle function (*51–53*). In this system, densely packed mitochondria exhibited improved calcium resilience and metabolic activity, suggesting a potential for spatial control over organelle function within engineered matrices. By measuring the MMP level, ATP production ability and the inner membrane cristae morphology of mitochondria collected after 1.8 mmol/L Ca^2+^ treatment, we conclude that the Mito@LLPS hydrogel has a good protective effect on mitochondria (*54*). The combined results from ICP-MS analysis, MMP measurements, and ATP production assays of aggregated mitochondria provide mechanistic insight into why the Mito@LLPS hydrogel offers superior protection against Ca^2+^-induced mitochondrial damage compared to traditional single-phase hydrogel (Mito@gelatin). The phase-separated structure of hydrogels endows the system with the ability of Ca^2+^ buffering via Ca^2+^ heterogenous distribution and mitochondria crowding effect to preserve mitochondrial integrity and function. Importantly, the Mito@LLPS hydrogel also significantly enhances the long-term mitochondrial activity at 4°C, thereby further expanding its potential for clinical translation and off-the-shelf therapeutic applications.

In addition, the Mito@LLPS hydrogel exhibits several advantageous therapeutic properties compared to traditional chemically cross-linked and single-phase hydrogels. These characteristics include its rapid degradability, efficient systemic clearance, and timely mitochondrial release, all of which contribute to enhanced therapeutic efficacy and biosafety. The presence of LLPS elevates the sol-gel transition temperature. Remarkably, Mito@LLPS hydrogel retains its gel status at body temperature once injected into the myocardial tissue. This property enhances both hydrogel and mitochondria retention within myocardial tissue, enabling sustained mitochondrial release over time. Besides, compared to covalently cross-linked hydrogels, the relatively poor mechanical stability of this physically cross-linked hydrogel facilitates efficient and timely mitochondrial release and uptake by damaged cardiomyocytes. Confocal microscopy reveals that labelled mitochondria are rapidly released from the hydrogel and internalized by H9c2 cardiomyocytes in vitro. Comparable mitochondria uptake is also observed in vivo following intramyocardial injection in rat models. Importantly, LLPS-assisted delivery results in prolonged retention of transplanted mitochondria in myocardial tissue and reduced systemic clearance compared to naked mitochondria, which are rapidly metabolized through liver and kidney pathways. Furthermore, Mito@LLPS hydrogel demonstrates excellent myocardial biocompatibility, preserves cardiac function, and elicits no observable toxicity in major organs. Collectively, these results confirm PEG/gelatin-based LLPS hydrogels as effective and biocompatible delivery vehicles for mitochondria transplantation. Moreover, the results of Seahorse energy metabolism analysis reveal that the OCR of cells subjected to H/R is significantly reduced, indicating a compromised capacity for aerobic respiration. In contrast, the ECAR is elevated, reflecting enhanced anaerobic glycolysis. This shift is associated with mitochondrial damage induced by H/R, resulting in disrupted cellular energy homeostasis and increased reliance on glycolysis for ATP production. However, ATP yield from glycolysis is markedly lower than OXPHOS, leading to an overall energy deficit and a reduced intracellular ATP level. Notably, cells treated with the Mito@LLPS hydrogel exhibit a superior recovery of energy metabolism following H/R injury compared to those treated with naked Mito or Mito@gelatin, likely owing to the enhanced mitochondria protective effects of the Mito@LLPS hydrogel.

Moreover, the internalization of exogenous mitochondria by recipient cells has previously been demonstrated to occur via several mechanisms, including endocytosis, membrane fusion, and tunnelling nanotubes (TNTs) (*55*). To investigate the fate of transplanted mitochondria via Mito@LLPS hydrogel following internalization, we conduct a series of molecular biology experiments to assess their impact on mitochondrial dynamics (*20, 56*). WB analysis reveals that H/R injury significantly suppresses the expression of mitochondrial fusion-related protein MFN1 (*57*), while elevating the expression of the fission-related protein FIS1 (*58*). Notably, mitochondria transplantation reverses this trend, leading to a significant upregulation of MFN1, although changes in FIS1 expression do not reach statistical significance. Consequently, the ratio of fusion to fission proteins in transplanted cells approached levels observed in normal cells, suggesting that mitochondria transplantation helps restore the dynamic balance between mitochondrial fusion and fission, thereby promoting mitochondrial homeostasis. Notably, this restorative effect is more pronounced in the Mito@LLPS hydrogel group than that in either the naked mitochondria or Mito@gelatin groups, emphasizing the positive value of the phase-separated structure of hydrogels in preserving mitochondrial integrity and function.

Furthermore, systematic in vivo evaluation in a rat of the MIRI model confirms the therapeutic efficacy of Mito@LLPS hydrogel. Notably, LLPS-mediated mitochondria transplantation significantly enhances cardiac functional recovery, as evidenced by the increase in LVEF and LVFS as measured via echocardiography. Additionally, this strategy significantly reduces myocardial fibrosis. These findings are consistent with the observed cellular and molecular benefits, collectively highlighting the superior efficacy of Mito@LLPS hydrogel over conventional mitochondria transplantation approaches via single-phase hydrogels.

This study also offers broader implications for mitochondria transplantation beyond myocardial applications. Given the role of mitochondrial dysfunction in a wide range of acute organ injuries, such as hepatic I/R injury, stroke and acute kidney I/R injury, the Mito@LLPS hydrogel platform may be adapted for organ-specific delivery of functional mitochondria. Furthermore, the phase-separated hydrogel design is inherently tunable, allowing modulation of mechanical properties, degradation rates, and cargo release kinetics to match specific therapeutic needs.

Despite these promising findings, several limitations should be acknowledged. First, although mitochondrial internalization and functional integration have been confirmed, the long-term fate of the transplanted mitochondria within recipient cells remains unclear. Critical questions persist regarding whether the exogenous mitochondria genetic material remains functionally active, whether it integrates into the host cell’s genome, and whether it is inherited by daughter cells during mitosis. Thus, further investigation is required to elucidate the long-term persistence, genomic integration, and mitotic transmission of transplanted mitochondria in recipient cells (*59*). Second, it is critical to study the response of other cell populations of the heart—such as resident macrophages, endothelial cells, and fibroblasts—to mitochondria transplantation. Specifically, whether exogenous mitochondria are taken up by these cell types within the infarcted region, and how such interactions influence their functional behavior, requires further comprehensive analysis.

In summary, the Mito@LLPS hydrogel we developed represents a novel biomaterial-based strategy for enhancing mitochondria transplantation efficiency by integrating physical protection, ionic buffering, and mitochondria crowding effect into a single delivery platform. By addressing key barriers associated with mitochondrial viability and internalization, this approach significantly enhances therapeutic effects in MIRI compared to conventional MTT via single-phase hydrogels and may serve as a blueprint for future organelle-based therapies across diverse clinical contexts.

## Materials and Methods

### Materials

Tissue Mitochondrial Isolation Kit (C3606), MitoTracker Green FM (C1048), Bradford Protein Assay Kit (P0006), Enhanced ATP assay kit (S0027), Hoechst 33342 stains (C1022), TMRE stains (C2001S), RIPA lysis buffer (P0013B), Blocking Buffer for Western Blot (P0252) and Primary Antibody Dilution Buffer for Western Blot (P0023A) were purchased from Beyotime Biotechnology (PRC). Polyethylene glycol (M_w_ = 10,000) was purchased from SINOPEG (PRC). Calcium chloride dihydrate (98%) was purchased from Biodepharm (PRC). DSPE-PEG-Rhodamine B (M_w_ = 10,000) was purchased from Tanshtech (PRC). Gelatin from porcine skin (39465), disodium succinate hexahydrate (BioReagent), osmium tetroxide (4 wt% in H_2_O), potassium ferrocyanide trihydrate (≥99.95%) and epoxy resin (≥95%) were purchased from Sigma-Aldrich (USA). Gelatin Methacryloyl (EFL-GM-30) was purchased from EFL (PRC). Glutaraldehyde (2.5%) was purchased from EcoTop Bio (PRC). 96-well cell culture plates (black frame, clear flat bottom, 3340) and Transwell upper compartment (6.5 mm Transwell® 8.0 µm, 3422) were purchased from Corning (USA). Adenosine diphosphate (≥95%) and uranyl acetate (99%) were purchased from Acmec (PRC). Sodium cacodylate buffer (0.15 mol/L, pH 8.5) was purchased from YuanYe (PRC). CCK8 (HY-K0301), JC-1 stain (CBIC2), MitoSOX Red (HY-D1055), MitoTracker Deep Red FM (HY-D1783) and DAPI dihydrochloride (HY-D0814) were purchased from MCE (USA). Calcein-AM/PI Double Staining Kit (110–116) was purchased from GOONIE (PRC). Seahorse XF cell mitochondrial stress assay kit (103015–100), Seahorse XF Real-time ATP Rate assay kit (103592–100) and Seahorse XF glycolysis rate assay kit (103344–100) was purchased from Agilent (USA). Antifade mounting medium (S2100), phenylmethylsulphonyl fluoride (PMSF, P8340) and iFlour 555-conjugated WGA (I3310) were purchased from Solarbio (PRC). One-step TUNEL In Situ Apoptosis Kit (Green, FITC) (E-CK-A320) was purchased from Elabscience (PRC). Anti-MFN1 (ab126575), anti-FIS1 (ab229969), anti-GAPDH (ab8245), Anti-alpha SMA (ab124964), Anti-Sarcomeric α-Actinin (ab9465), anti-rabbit HRP-conjugated IgG (ab6721), anti-mouse HRP-conjugated IgG (ab6728) and Goat Anti-Mouse IgG H&L (Alexa Fluor® 488) (ab150113) were purchased from Abcam (USA). ECL detection reagent (P10060) was purchased from NCM Biotech (PRC).

### Animals use and study approval

Wild-type (WT) Sprague-Dawley (SD) rats (240-280 g; male; 8 weeks old) were procured from Laboratory Animal Center, Southern Medical University (Guangzhou, PRC). The animals were randomly grouped for treated and untreated controls, and sacrificed by carbon dioxide anesthesia. All experiments and animal procedures were sanctioned by the Ethics Committee of the Tenth Affiliated Hospital of Southern Medical University (Dongguan People’s Hospital) (IACUC-AWEC-202302015) and carried out following the Regulations for the Administration of Affairs Concerning Experimental Animals (P. R. China).

### Cell culture

The H9c2 cell line was purchased from Guangzhou Xinyuan Electronic Technology Co., Ltd. and maintained in Dulbecco’s modified Eagle medium (DMEM, high glucose, 1.5% NaHCO_3_) supplemented with 10% fetal bovine serum (FBS) and 1% penicillin-streptomycin amphotericin B solution. Cells were placed into humidified incubator with 5% CO_2_ and at 37 °C. No mycoplasma and bacterial contamination were detected in the H9c2 cell line.

### Isolation and quantification of mitochondria

Mitochondria were isolated from the heart tissue of wild-type SD rats using a mitochondrial isolation kit following the manufacturer’s protocol. All operations were conducted in an ice bath. The isolated mitochondria were labelled with MitoTracker Green FM (100 nmol/L), and the number of mitochondria was determined by flow cytometry (NovoCyte Quanteon, Agilent, USA). Specifically, the volume corresponding to 10,000 events was measured by flow cytometry to calculate the mitochondrial concentration in the suspension. An aliquot of the mitochondrial suspension was lysed with mitochondrial lysis buffer to extract mitochondrial proteins. Bradford Protein Assay Kit was used to measure the protein concentration according to the kit’s instructions. Subsequently, the number of mitochondria was estimated based on the approximation of the quantity of mitochondrial proteins.

### Preparation of LLPS hydrogel, naked Mito, Mito@gelatin and Mito@LLPS hydrogel

The Polyethylene glycol (PEG) and gelatin powder were sterilized under an ultraviolet lamp for 10 min. For solution preparation, 0.5 g of PEG powder were dissolved in 2.0 mL of physiological saline or PBS and then stirred at RT until the solution turned clear. 2.0 g of gelatin powder were dissolved in 9 mL of physiological saline or PBS and then heated at 45 °C with stirring until dissolved. Then the dissolved PEG solution and gelatin solution were gently mixed by pipetting to obtain PEG/gelatin liquid-liquid phase separated hydrogel. LLPS hydrogel: the pre-prepared PEG/gelatin LLPS hydrogel. Naked Mito: 0.1 mL of freshly isolated mitochondria were resuspended in 0.9 mL of PBS. Mito@gelatin hydrogel: 0.1 mL of freshly isolated mitochondria were resuspended in 0.9 mL of pre-prepared gelatin (18.2 wt%) solution. Mito@LLPS hydrogel: 0.1 mL of freshly isolated mitochondria were resuspended in 0.9 mL of pre-prepared PEG/gelatin LLPS hydrogel.

### Mito@LLPS hydrogel confocal microscopy imaging

The continuous and dispersed phases of phase separation hydrogels were determined by mixing gelatin solution (contained 5% gelatin-FITC) and PEG solution (contained 5% DSPE-PEG-Rhodamine), followed by gentle pipetting. MitoTracker Green FM (100 nmol/L) labelled freshly isolated mitochondria were used to determine their distribution within the phases. LCSM (Zeiss LSM900 with Airyscan2, GER) was used to image.

### Scanning electron microscope image

2.5% glutaraldehyde was used to fix freshly isolated mitochondria at 4 °C overnight, and then washed with PBS or normal saline. Subsequently, the mitochondria samples were dehydrated through a graded ethanol series (30%, 50%, 70%, 80%, and 90%), Prepared by mixing anhydrous ethanol and ddH₂O. This was followed by two treatments with absolute ethanol for 15 min each. Finally, the samples were thoroughly dried. The Mito@LLPS sample was obtained by mixing the dehydrated mitochondrial samples with PEG/gelatin LLPS hydrogels. The prepared PEG, gelatin, Mito@LLPS hydrogel were lyophilized overnight in a vacuum lyophilizer (Christ/Alpha 1-2 LD plus, GER) and then transferred to the sample stage and used Ion Sputter to spray with gold for 2 min. Apreo 2 SEM was used to image.

### Rheological properties

The rheological properties of the gelatin and PEG/gelatin LLPS hydrogel with different concentrations were analyzed using a dynamic stress-controlled rheometer (Anton Paar MCR102e, ATU) equipped with cone plate (gap: 1 mm, diameter: 60 mm). Multiple different detection methods were used to characterize the mechanical properties of the LLPS hydrogels and single-phase hydrogel. The rheological properties of the hydrogels were measured by the temperature scanning method within a temperature range of 20-50 °C.

### Detection of Ca^2+^ distribution coefficient 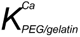

The distribution coefficient 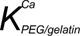 is defined as the ratio of the concentration of Ca^2+^ in the gelatin phase (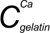) to the concentration of (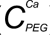) in the PEG phase:

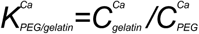

Hence, the value of 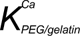 greater than 1 indicated a higher calcium ion concentration in the gelatin phase than in the PEG phase. 1.8 mmol/L Ca^2+^ solution was prepared using calcium chloride (CaCl_2_) and ddH_2_O. This Ca^2+^ solution was used to separate the PEG and gelatin phases by centrifugation.

PEG and gelatin solutions were mixed and blown well in proportion, then immediately performed a centrifugation at 12,000×g for 10 min. After delamination of the two-phase solution, the upper PEG solution was transferred to other Eppendorf tubes. Their densities were calculated by injecting known volumes of PEG and gelatin solutions through a syringe pump and weighing them, because these solutions were too viscous to accurately use a pipette to draw an accurate volume. Subsequently, appropriate amounts of gelatin and PEG solutions were accurately weighed and mixed. Pure nitric acid was then added to the mixture, and then incubated overnight in a 70 °C metal bath. Then the samples were diluted using ddH_2_O and then filtered through 0.22 μm filter membrane. iCAP™ TQ ICP-MS (Thermo Scientific, USA) was used to detect Ca^2+^ content. The distribution coefficient of Ca^2+^ was calculated.

### Mitochondrial membrane potential (MMP) detection

Freshly isolated mitochondria were stained with JC-1 (5 μg/mL) at 4 °C for 10 min and washed with mitochondrial storage buffer (MSB) for 2 times. The labelled mitochondria were then divided into the four groups (control group, naked Mito group, Mito@gelatin group and Mito@LLPS hydrogel group) and resuspended in PBS or hydrogel to 100 μL and placed into 96-well cell culture plates (black frame, clear flat bottom). Subsequently, 100 μL of Hank’s Balanced Salt Solution (HBSS) containing 1.8 mmol/L Ca^2+^ was added to each well. The intensity of fluorescence (excitation/emission, 485 nm/590 nm) was time-scanned in one-hour intervals, with checks conducted every 5 min by multifunctional microplate reader.

Moreover, the MMP of isolated mitochondria which were suspended in PEG in different concentrations (4×10^4^ particles/mL, 8×10^4^ particles/mL, 1.6×10^5^ particles/mL) were also measured using the similar method, separately.

Furthermore, the MMP of isolated mitochondria which were store at 4 °C for 0, 4, 8 h were also measured using the similar method, separately.

Moreover, the MMP of isolated mitochondria which were suspended in PEG in different concentrations (4×10^4^ particles/mL, 8×10^4^ particles/mL, 1.6×10^5^ particles/mL) were also measured using the similar method, separately.

Furthermore, the MMP of isolated mitochondria which were store at 4 °C for 0, 4, 8 h were also measured using the similar method, separately.

### Measurement of isolated mitochondrial ATP production level

Experimental groups (control, naked Mito, Mito@gelatin and Mito@LLPS hydrogel) respectively resuspended in PBS, PEG, gelatin, and PEG/gelatin LLPS hydrogel to a volume of 200 μL and placed into 24-well cell culture plates. Subsequently, 200 µL of HBSS buffer was added and incubated at 37 °C. After a half-hour incubation, 200 μL of HBSS containing 75 mmol/L succinate and 4.95 mmol/L ADP was added, then incubated for 30 min at 37 °C. The solution was then placed to a 1.5 mL tube, centrifuged at 12,000×g for 10 min at 4 °C. 100 μL ATP assay working solution was then dispensed into black-walled, clear flat-bottom 96-well cell culture plates. Following a 5-minute equilibration period at RT, 20 μL of either cell supernatant or ATP standard was added. Chemiluminescence was measured by multifunctional microplate reader. Moreover, the ATP production level of isolated mitochondria which were suspended in PEG in different concentrations (4×10^4^ particles/mL, 8×10^4^ particles/mL, 1.6×10^5^ particles/mL) were also measured in the similar method, separately.

Furthermore, the ATP production level of isolated mitochondria which were store at 4 °C for 0, 4, 8 h were also measured in the similar method, separately.

### Transmission electron microscopy image

The experiment was conducted to four groups (control, naked Mito, Mito@gelatin and Mito@LLPS hydrogel). In the control group, freshly isolated mitochondrial were immediately immobilized with 2.5% glutaraldehyde for 1 h on ice. The remaining three groups were suspended in PBS buffer, gelatin, and PEG/gelatin hydrogel, respectively. An equal volume of HBSS buffer added to each suspension, followed incubating at 37 °C for 1 h. Subsequently, a large volume of 2.5% glutaraldehyde was added to each group for fixation on ice for 1 hour. Afterwards, the samples were rinsed in a 0.15 mol/L sodium cacodylate buffer containing 3 mmol/L of CaCl_2_ (in an ice-water environment) to maintain the integrity of the cell membrane preservation. The fixation treatment was completed using a 1% osmium tetroxide solution in 0.15 mol/L sodium cacodylate sulfonate buffer (containing 0.8% potassium ferrocyanide and 3 mmol/L CaCl_2_) in an ice-water environment. After repeated rinsing in ice water, the mitochondria were stained, and the inner membrane was stabilized in a 2% acetic uranyl solution in ice water. Dehydration process was carried out successively with 30%, 50%, 70% and 90% ethanol gradient solutions on ice. Dehydration was completed at RT with 100% ethanol. Infiltration was performed with a homogeneously mixed 1:1 (v/v) solution of ethanol and epoxy resin. The infiltration process continued, and 100% epoxy resin exchanges were carried out multiple times. Thin slices (with a thickness of 80 nanometers) cut from the sample were prepared using an ultrathin sectioning machine (Leica EM UC7, Germany). After post-staining with uranyl acetate and lead salt, these sections were observed using a TEM (Tauros L120C from Thermo Fisher Scientific, USA). The images were recorded at a magnification of 12,000×.

### Mitochondrial Release Assay

The suspension containing freshly isolated mitochondrial were divided equally into four groups: control, naked Mito, Mito@gelatin and Mito@LLPS hydrogel. Mitochondria in the remaining three experimental groups were resuspended and thoroughly mixed with PBS, gelatin, or LLPS hydrogel, respectively. Following centrifugating at 12,000×g for 10 min at 4 °C, 100 μL resulting solution was carefully transferred to the upper compartment of a Transwell system. In addition, an appropriate amount of medium (without cells) was placed into the lower compartment, and then placed into incubator. Subsequently, mitochondria in the culture medium of the lower compartment were collected after incubating 1 h by centrifugating at 12,000×g for 5 min at 4 °C. Mitochondrial proteins were then extracted to determine the amounts of mitochondria.

### In vivo distribution of transplanted mitochondria

Freshly isolated mitochondria were labelled with MitoTracker Deep Red FM (100 nmol/L). The labelled mitochondria were then mixed with PBS and LLPS hydrogel, respectively. 100 μL of solutions (naked Mito, Mito@LLPS hydrogel) were injected into each SD rat through intramyocardial injection. SD rats were sacrificed by carbon dioxide anesthesia at 6 h, 12 h, 24 h, and 48 h after post-intramyocardial injection, and major organs (heart, liver, spleen, lung, kidney) were collected for IVIS fluorescence imaging in vitro. The fluorescence intensity of each organ was photographed by PerkinElmer IVIS Spectrum (PerkinElmer, USA, Luminescent exposure: 2.5 sec, f: 2.0, excitation/emission; 605 nm/660 nm).

### Assessment of transplanted mitochondria uptake in vivo

The MitoTracker Deep Red FM (100 nmol/L) labelled isolated mitochondria were condensed in PEG/gelatin LLPS hydrogel (1×10^7^ particles/mL). Then, 100 μL of Mito@LLPS hydrogel was injected into the left ventricle. SD rats were euthanized by carbon dioxide inhalation at 6 h after intramyocardial injection. The heart tissue samples were embedded and sectioned by freezing microtome. 4% paraformaldehyde was used to fix the sections at RT. Subsequently, anti-Sarcomeric α-Actinin and DAPI dihydrochloride solution (10 μg/mL) was applied to each section and incubated at RT for 30 min. Finally, antifade mounting medium was applied to seal the slides, and imaging was performed using LSCM.

### Assessment of transplanted mitochondria uptake in vitro

H9c2 cells were seeded onto round coverslip (Φ 14 mm) in 24-well cell culture plate. MitoTracker Green FM (100 nmol/L) labelled freshly isolated mitochondria were suspended with PBS, gelatin and PEG/gelatin LLPS hydrogel (5×10^6^ particles/ml). Then, 100 μL of naked Mito, Mito@gelatin and Mito@LLPS hydrogel were placed into Transwell upper compartment and co-incubated with H9c2 cells in 24-well cell culture plate for 6 h. Then, H9c2 cell line stained with Hoechst 33342 (5 μg/mL). Laser Scanning confocal microscope was used to image.

### Cellular hypoxia-reoxygenation model

H9c2 cells were seeded into a cell plate. The following day, the medium was removed, and replaced with serum-free DMEM (without glucose, HEPES or sodium pyruvate). Subsequently, the plate was transferred to an anaerobic cell culture incubator (ESCO, CCL-170T-8, Singapore) maintained at 37 °C under a controlled atmosphere of 5% CO₂ and 1% O₂. After anaerobic incubation for 6 h, the cells were removed from the anaerobic chamber, washed with PBS, and resuspended in fresh DMEM. They were then transferred to a standard cell culture incubator for an additional 6 h under normoxic conditions.

### Cellular relative active assay

The cell suspension (2.5×10^6^ cells/well, 0.5 mL/well) was seeded into 24-well cell plate to incubate overnight. Following H/R treatment, naked Mito, Mito@gelatin and Mito@LLPS hydrogel were added to the Transwell upper compartment, and incubated with the treated cells for 24 h. Both culture medium and the contents of the Transwell upper compartment were carefully removed, followed by gentle washing with PBS. Subsequently, 0.45 mL of culture medium supplemented with 50 μL CCK-8 solution was added and then incubated for 1 hour under standard conditions. Then, 100 μL supernatant was transferred to a 96-well plate for subsequent analysis. The absorbance was measured at 450 nm by a multifunctional microplate reader.

For safety evaluation of hydrogels in vitro, Cell suspension (6,000 cells/well, 100 μL/well) was inoculated into a 96-well plate. Subsequently, after cell adhesion, mixed 90 μL culture medium with 10 μL of PBS or LLPS hydrogel, was added to every well and incubated for 24 h, cytotoxicity of the LLPS hydrogel was detected by CCK-8 assay.

### Detection of MMP (ΔΨm) and mitochondrial superoxide

The cell suspension (5×10^6^ cells/well, 2 mL/well) was seeded into a 6-well culture plate. Following H/R treatment, PBS, naked Mito, Mito@gelatin and Mito@LLPS hydrogel were added to the Transwell upper compartment, and incubated with the treated cells for 24 h. Cells were digested and stained using 1 mL TMRE and 1 mL MitoSOX Red (5 μmol/L) staining work solution. After incubation for 20 min, the cells were precipitated by centrifugation at 600×g and 4 °C for 3 min. The fluorescence was analyzed by flow cytometry.

### Seahorse cell energy and metabolism measurement

OCR was measured using the Seahorse XF cell mitochondrial stress assay kit. Cells were seeded into XFe96/XF Pro Cell Culture Microplate at 8,000 cells/well and incubated overnight. After establishing the hypoxia-reoxygenation (H/R) model, cells were treated with PBS, naked Mito, Mito@gelatin and Mito@LLPS hydrogel, respectively. Following treatment, the cell culture medium was removed, cells were rinsed once with PBS, and replaced with Seahorse XF assay medium. This medium consisted of Seahorse XF DMEM medium, supplemented with 10 mmol/L glucose, 1 mmol/L pyruvate and 2 mmol/L glutamine. OCR analyses were conducted under basal conditions and following the subsequent injections of oligomycin (1.5 μmol/L), carbonyl cyanide-4-(trifluoromethoxy) phenylhydrazone (FCCP, 2 μmol/L) and rotenone plus antimycin A mix (Rot/AA, 0.5 μmol/L) with Hoechst 33342 (5 μg/mL) as a pilot experiment. These procedures were meticulously executed according to the protocol.

The ATP real-time production rate was quantified using Seahorse XF Real-time ATP Rate assay kit. The cell processing protocol and DMEM composition were identical to those employed in the OCR detection assay, with the sole variation being the drug treatment. Specifically, oligomycin (1.5 μmol/L) and rotenone/antimycin A (Rot/AA, 0.5 μmol/L) were administered sequentially, along with Hoechst 33342 (5 μg/mL), as indicated.

### Western Blot analysis

Cell samples were lysed with RIPA lysis buffer (supplemented with 1 mmol/L phenylmethylsulphonyl fluoride) and scraped and transferred into 1.5 mL tubes. The protein supernatant was taken by centrifugation at 12,000×g at 4 °C for 5 min. Bradford assay kit was used to determine the protein concentration. The protein samples were then mixed with 5× SDA-PAGE loading buffer (with DTT) and heated at 95 °C for 10 min. The samples were run on a 7.5% SDS-PAGE polyacrylamide gel and transferred onto a polyvinylidene difluoride (PVDF) membrane. The membrane was blocked using a blocking buffer and subsequently incubated with primary antibody, including anti-MFN1 (1:1,000), anti-FIS1 (1:1,000), anti-GAPDH (1:1,000), and ATPB (1:1,000), all diluted in a primary antibody dilution buffer. Following primary antibody incubation, membranes were treated with either HRP-conjugated anti-rabbit IgG (1:5,000) or HRP-conjugated anti-mouse IgG (1:5,000) as appropriate. Protein signals were then visualized using an ECL detection reagent. The nearest molecular weight markers (10-200 kDa) were shown to the left of the blots in all figures. WB images were analyzed using ImageJ software.

### ATP Assay of cellular and tissue

H9c2 cells were seeded into 6 cm cell culture plate. Following H/R treatment, naked Mito, Mito@gelatin, and Mito@LLPS hydrogel were added, and incubated for 24 h. The tissue samples obtained from rats in the MIRI model at seven days after operation and the treatment of LLPS hydrogel, naked Mito, Mito@gelatin, and Mito@LLPS hydrogel, and then 20 mg of tissue samples were collected from the injured area. Then the samples were homogenized in an appropriate volume of lysis buffer. After that, the homogenate was placed in 1.5 mL tubes and centrifuged at 12,000×g for 5 min at 4 °C to obtain the cell supernatant. Subsequently, 100 μL of the ATP assay working solution was added to each well. Following a 5-minute equilibration period at RT, 20 μL of cell supernatant was introduced into the assay well. The detection of chemiluminescence was carried out using a multifunctional microplate reader. Protein concentrations were detected by Bradford protein assay kit. ATP levels were normalized to protein concentration, and the final concentrations of ATP were converted to nmol/mg of protein.

### The establishment of myocardial I/R model and intramyocardial injection of mitochondria samples

Surgical induction of cardiac ischemia/reperfusion was performed on SD rats (male, about 240-280 g, 8 weeks old) according to the previous report. Briefly, all rats were anaesthetized by 2% isoflurane. A surgical incision was made to open the third intercostal space above the left thorax, allowing exposure of the heart. The pericardium was carefully incised to reveal the cardiac surface. The branches of the left coronary artery extend downward and are arranged parallel to the tip of the left atrial appendage, which was identified and ligated using a 6-0 suture. The observation of white color of myocardial tissue below the ligation line indicated that the operation was successful. 45 min after ischemia, the slip knot was released by using microscopic scissors to cut open the knot to reperfuse the myocardium. Immediately after reperfusion, 100 μL of solution (PBS, LLPS hydrogel, naked Mito, Mito@gelatin or Mito@LLPS hydrogel) was injected into the infarcted myocardium using a 30 G needle. Rats in the I/R group underwent ischemia/reperfusion injury followed by PBS injection, whereas control animals were subjected only to thoracotomy and PBS injection without ischemia/reperfusion surgery. Upon completion of the procedures, the incision was closed with layered sutures, the skin was disinfected using iodophor, and the animals were allowed to recover while supported on a ventilator. At day 28 after surgery, animals were sacrificed, and cardiac tissue was collected.

### Echocardiography

Induction of anesthesia was performed in rats with 2-3% isoflurane. Continuous administration of 1-1.5% isoflurane was maintained throughout the echocardiography procedure, ensuring a stable heart rate of approximately 350-500 beats per minute and a controlled body temperature of 37 °C. The M-mode echocardiography was acquired in the short axis by VisualSonics Vevo® 3100 Imaging System with an MX250 transducer (Fujifilm, JPN). The cardiac function measurements analyses were calculated by Vevo LAB 3.1.1 software.

### Masson’s trichrome staining

Tissue sections were stained with Masson’s trichrome stain. After staining, sections were rinsed under running water for 3 min, followed by dehydration through a graded ethanol series. Subsequently, the sections were cleared with xylene to achieve transparency and then mounted with neutral resinous mounting medium to cover the slides. The samples were scanned with the Slice Scanner System (Olympus-VS200 SLIDEVIE, JPN). Fibrosis was quantified by Fiji (ImageJ) software.

### Immunofluorescence staining

The tissue sections were stained with TUNEL apoptosis kit (Green, FITC), iFlour 555-conjugated WGA (10 μg/mL), Anti-alpha SMA primary antibody (diluted in 1:350) and Goat Anti-Mouse IgG H&L (Alexa Fluor® 488) (diluted in 1:3000). LSCM and Olympus Slice Scanner System were used to image. The number of apoptotic cells and the cross-sectional area of cardiomyocytes were measured using the Fiji (ImageJ) software

### Statistics

GraphPad Prism 8.0.2 (provided by GraphPad Company) were performed for graphs and statistical analyses. The statistical test methods employed are all explained in the legends.

## Supplementary Materials

This PDF file includes: Supplementary methods and Figs. S1 to S14

## Author contributions

Conceptualization: Z.L., X.W. and J.A. Methodology: J.A., Y.X., Q.L., Y.X., W.L., J.D., W.D., R.Z., S.M. and Y.Z. Investigation: X.W., J.A. and Y.X. Visualization: J.A. and Y.X. Supervision: Z.L. and X.W. Writing—original draft: X.W. and J.A. Writing—review & editing: Z.L., X.W., J.A., Y.X., Q.L., W.L. and J.D.

## Data and materials availability

All data needed to evaluate the conclusions in the paper are present in the paper and/or the Supplementary Materials.

