## Supporting information for "Transplantation of active mitochondria condensed in liquid−liquid phase-separated hydrogels ameliorates myocardial ischemia-reperfusion injury"

Jiacong Ai *et al.*

**This PDF file includes:**

Figs. S1 to S14

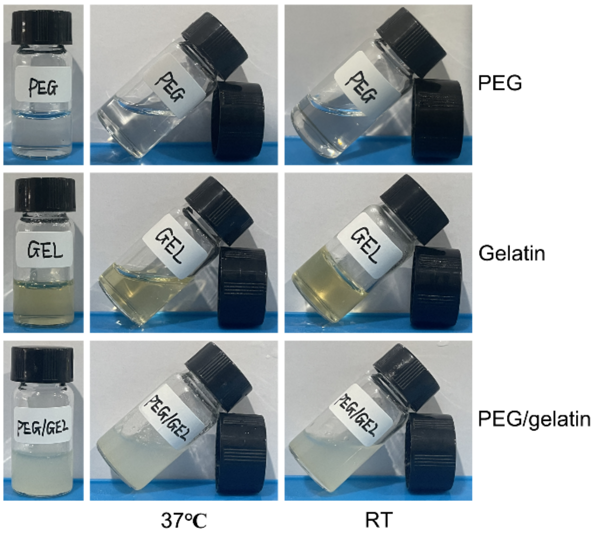


**Fig. S1. Photographs of PEG, gelatin, PEG/gelation solutions at 37 °C and room temperature (RT).** PEG (33.3 wt%) solution is a colorless, transparent liquid at both RT and 37 °C. In contrast, gelatin (18.2 wt%) exhibits thermo-sensitive behavior. It forms a yellow, non-flowing gel at RT but transitions into a yellow, flowable liquid when heated to 37 °C. PEG/gelatin mixture displays a homogeneous, cloudy opalescent appearance and maintains excellent fluidity under both 37°C and RT conditions.

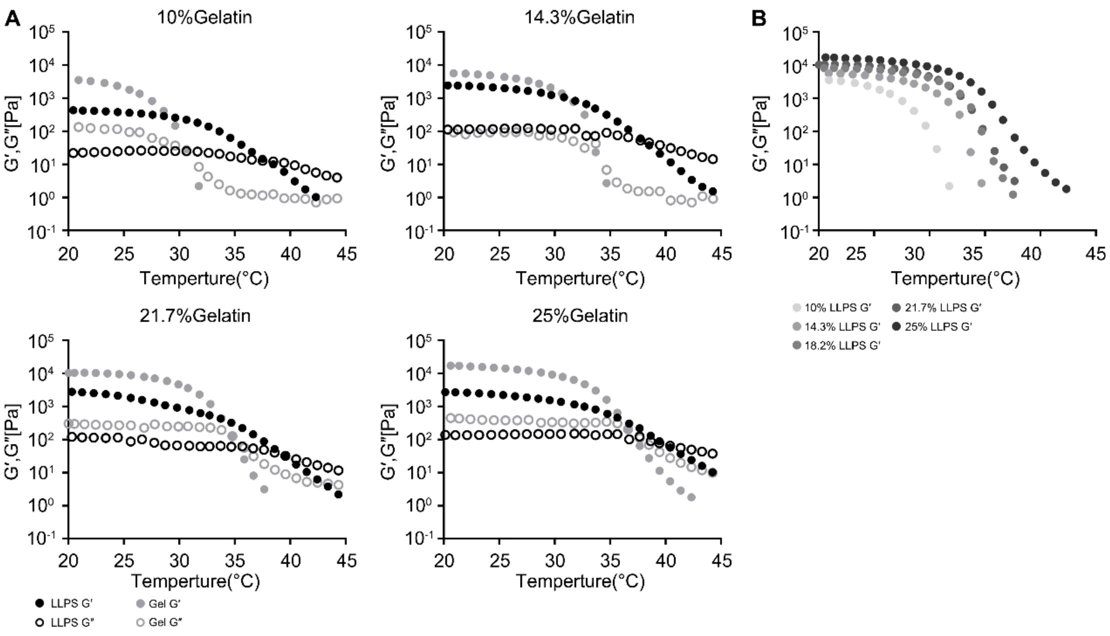


**Fig. S2. The rheological mechanics of pure gelatin and PEG/gelatin LLPS hydrogel under different concentrations of gelatin.** A, B. Detection of rheological mechanics (G', G") of PEG/gelatin LLPS hydrogel at different concentrations of gelatin (10, 14.3, 18.2, 21.7, 25 wt%) under various temperature conditions. The incorporation of PEG solution leads to an elevation in the sol-gel transition temperature of the PEG/gelatin LLPS hydrogel. Furthermore, a progressive increase in gelatin concentration results in a corresponding rise in the sol-gel transition temperature of the phase-separated hydrogel system.

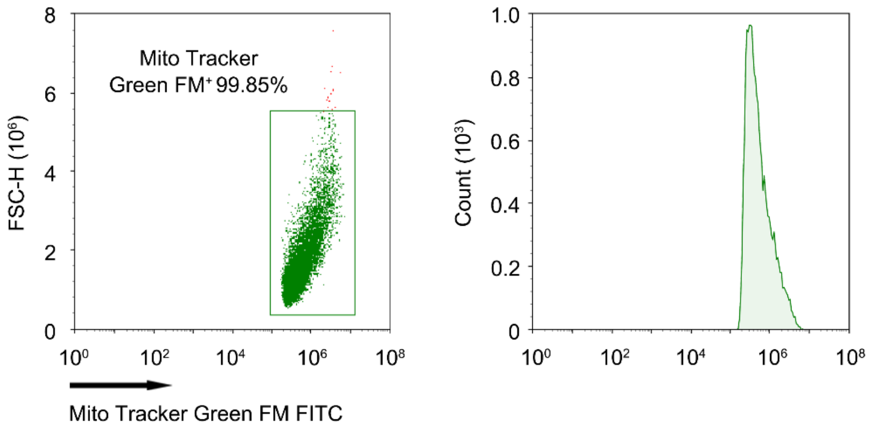


**Fig. S3. Flow cytometry of isolated mitochondria.** Freshly isolated mitochondria were stained with Mito Tracker Green FM and analyzed by flow cytometry. The results showed that nearly all mitochondria were successfully labelled, indicating that the isolation protocol yielded mitochondria with high purity.

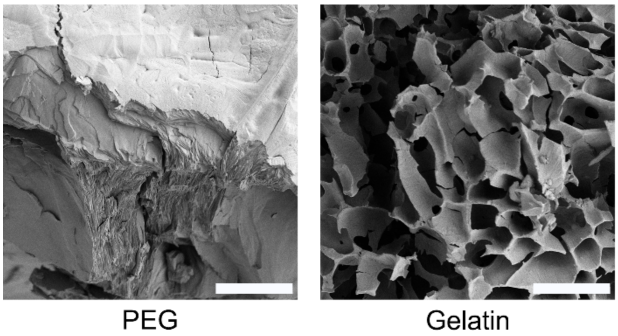


**Fig. S4. SEM images of PEG and gelatin solution.** SEM image of PEG solution (33.3 wt%) showed a dense and pore-free structure, which was similar to the structure of the PEG powder. SEM image of gelatin solution (18.2 wt%) showed a loose and porous structure. Scale bar, 10 μm.

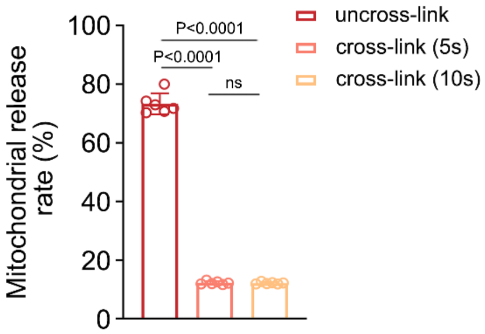


**Fig. S5. Mitochondrial release rate from uncross-linked and chemically cross-linked Mito@LLPS hydrogels.** Mito@LLPS hydrogels were subjected to UV-induced cross-linking using gelatin methacryloyl (GelMA) for 5 and 10 seconds, respectively. The influence of chemical cross-linking on mitochondrial release was systematically evaluated by comparing cross-linked and uncross-linked samples. Notably, chemical cross-linking significantly suppressed mitochondrial release, with release rates of 12.23% and 12.17% observed for the 5-second and 10-second cross-linked groups, respectively. In contrast, the uncross-linked group exhibited a markedly higher release rate of 66.32%, underscoring the effectiveness of GelMA cross-linking in stabilizing the hydrogel network and limiting cargo diffusion. *n*=6. Statistical significance was calculated via two-tail one-way ANOVA-Tukey's multiple comparisons test.

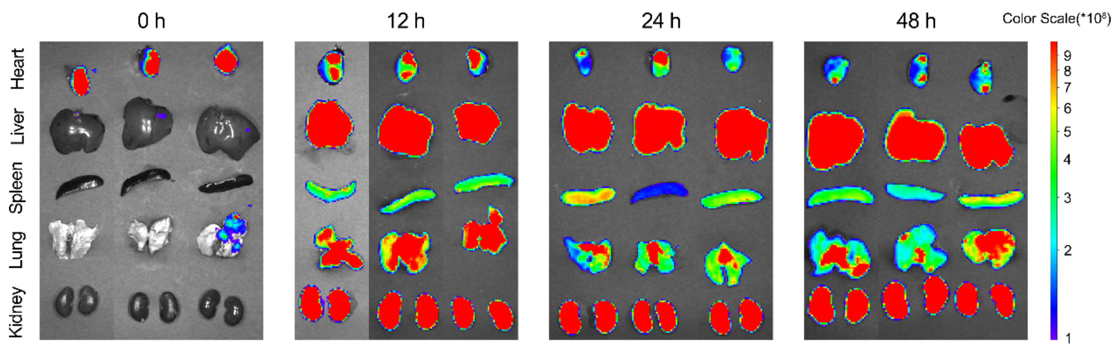


**Fig. S6. IVIS images of major organs of SD rats with intramyocardial injection of Mito@LLPS hydrogel.** To evaluate the in vivo distribution of the hydrogel system, Cy5-labelled gelatin was mixed with unlabeled gelatin to prepare the Mito@LLPS hydrogel. A total of 100 μL of the Mito@LLPS hydrogel was injected directly into the rat myocardium. At 0, 12, 24, and 48 hours post-intramyocardial injection, major organs—including the heart, liver, spleen, lung, and kidneys—were harvested for ex vivo imaging using IVIS.

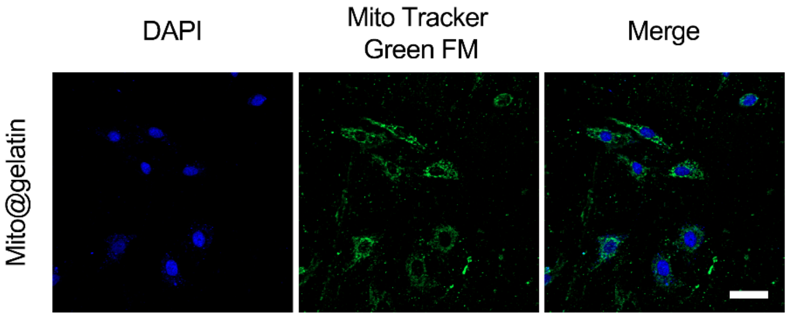


**Fig. S7. LCSM images of cellular uptake of Mito@gelatin.** H9c2 cells were co-incubated with Mito@gelatin using a Transwell system for 6 h. Cellular nuclei were stained with DAPI (blue), and transplanted mitochondria were visualized with MitoTracker Green (green). Scale bar, 20 μm.

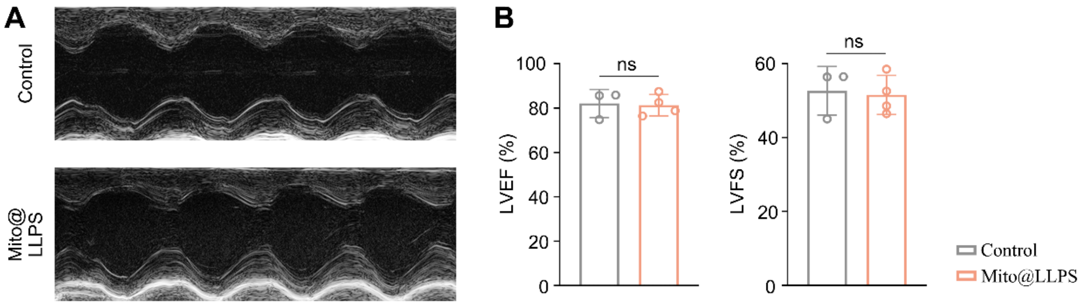


**Fig. S8. In vivo biosafety and biocompatibility of Mito@LLPS hydrogels.** 6-weeks SD rats were intramyocardially injected with 100 μL PBS or LLPS hydrogel. The cardiac ultrasound was used to evaluate the cardiac function indicators LVEF and LVFS of the SD rats 7 days after the operation. The results showed that the intramyocardial injection of the LLPS hydrogel did not affect the cardiac function level, suggesting that the LLPS hydrogel had good biosafety and biocompatibility. It also indicated that intramyocardial injection of Mito@LLPS into healthy SD rats did not promote their cardiac functions. Control group, $n$=3; Mito@LLPS group, $n$=4. Statistical significance was calculated via two-tail unpaired t-test.

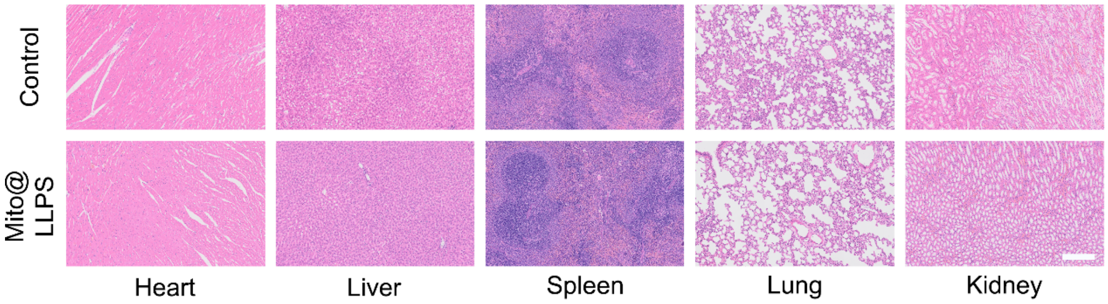


**Fig. S9. H&E staining images of the main organ tissue sections.** Main organ tissue H&E staining images of the SD rats at 28 days after injecting with 100 μL PBS (control group) or Mito@LLPS hydrogel. Scale bar: 250 μm.

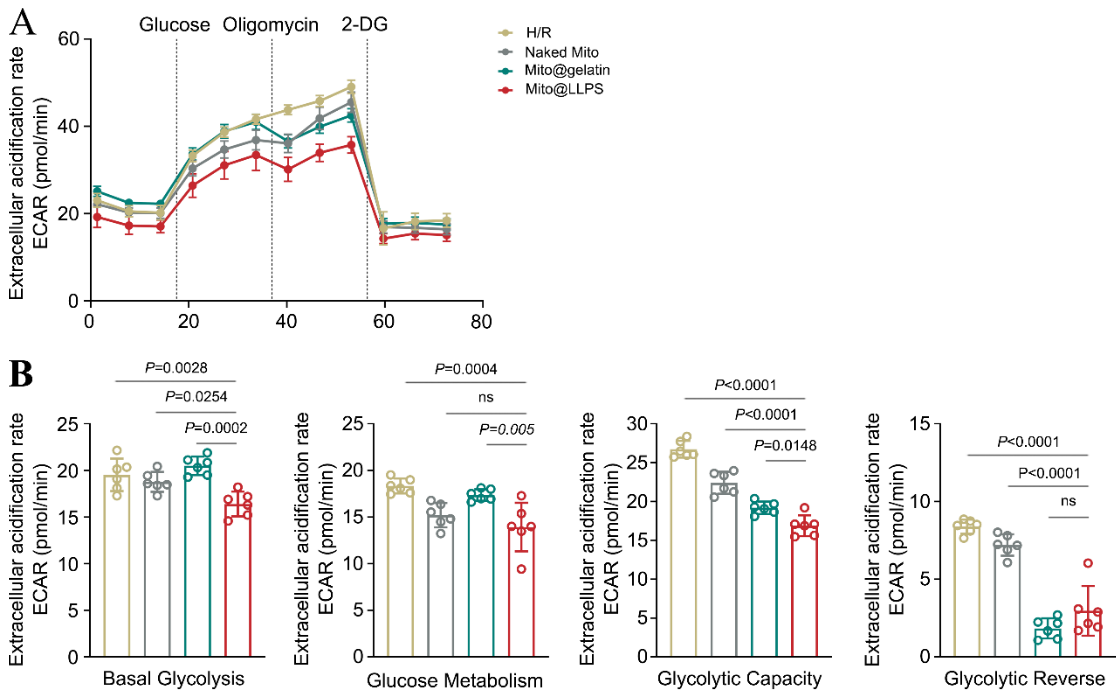


**Fig. S10. Results of cellular ECAR and quantitative analysis.** The results of cellular ECAR and quantitative analysis of basal glycolytic, glucose metabolism, glycolytic capacity and glycolytic reverse after MTT treatment of H/R-injured cells. *n*=6. Statistical significance was calculated via two-tail one-way ANOVA-Tukey's multiple comparisons test.

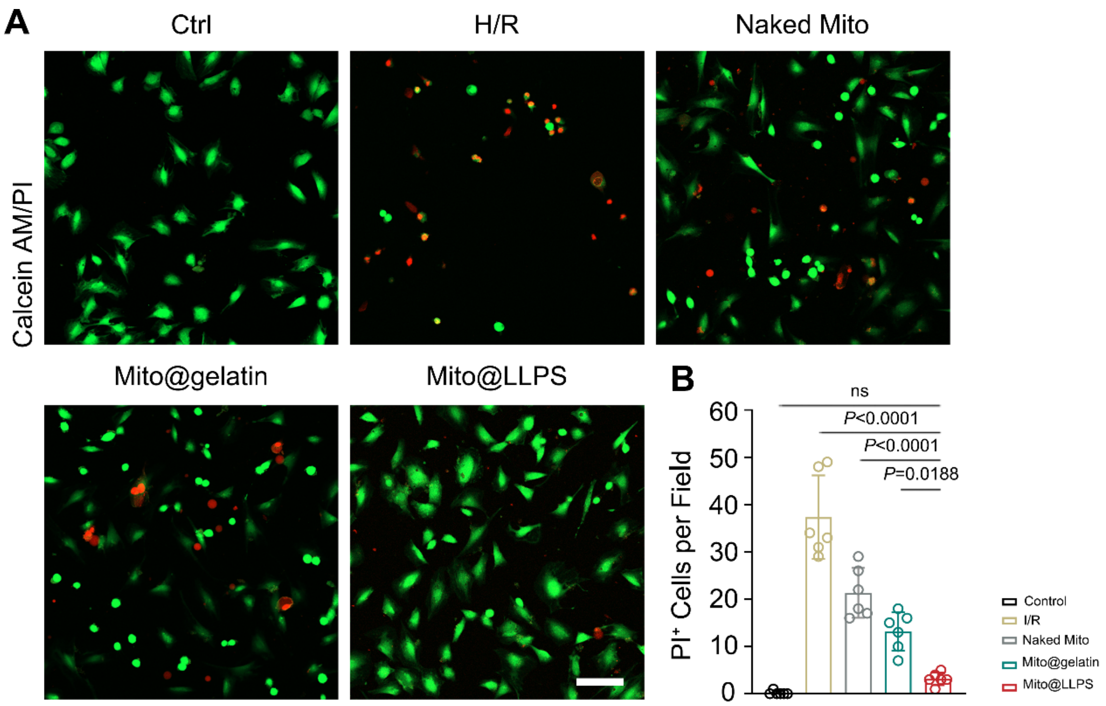


**Fig. S11. The LCSM images of cellular Calcein AM/PI double staining and quantitative analysis.** The LCSM images of cellular Calcein AM/PI double staining and quantitative analysis after MTT treatment of HR-injured cells. $n$=6. Statistical significance was calculated via two-tail one-way ANOVA-Tukey's multiple comparisons test. Scar bar: 20 μm.

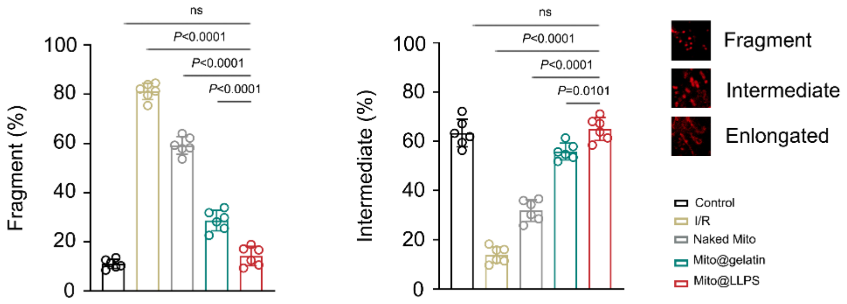


**Fig. S12. Quantitative analysis of mitochondria network morphology.** Quantitative analysis of mitochondria network morphology (fragment and intermediate) after different MTT treatments in H/R-injured cardiomyocytes (*n*=6). Statistical analysis was performed via two-tail one-way ANOVA-Tukey's multiple comparisons test.

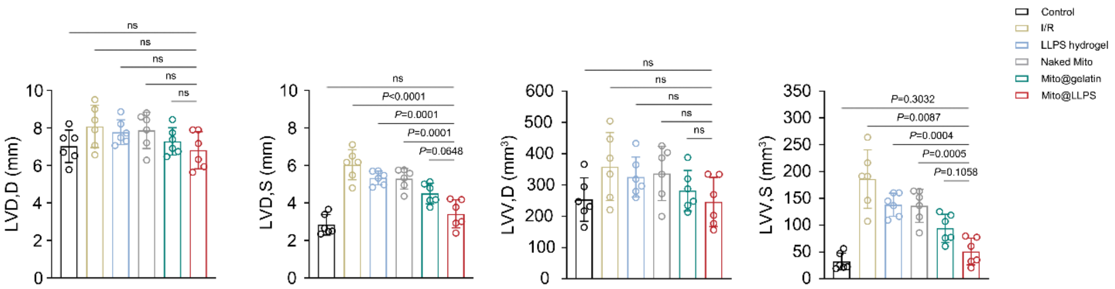


**Fig. S13. Quantitative analysis of echocardiography results.** The quantitative analysis of the echocardiography results of the SD rats on day 28 after surgery (*n*=6). Statistical significance was calculated via two-tail one-way ANOVA-Tukey's multiple comparisons test.

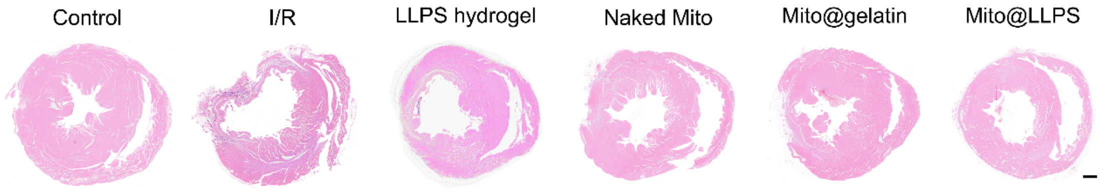


**Fig. S14. H&E staining of the heart tissue sections.** The images of H&E staining of the SD rat heart tissue sections on day 28 after surgery. Scale bar, 1 mm.
